# The regulation of Insulin/IGF-1 signaling by miR-142-3p associated with human longevity

**DOI:** 10.1101/2023.05.19.541542

**Authors:** Xifan Wang, Hwa Jin Jung, Brandon Milholland, Jinhua Cui, Cagdas Tazearslan, Gil Atzmon, Xizhe Wang, Jiping Yang, Qinghua Guo, Nir Barzilai, Paul D. Robbins, Yousin Suh

## Abstract

MicroRNAs (miRNAs) have been demonstrated to modulate life span in the invertebrates *C. elegans* and *Drosophila* by targeting conserved pathways of aging, such as insulin/IGF-1 signaling (IIS). However, a role for miRNAs in modulating human longevity has not been fully explored. Here we investigated novel roles of miRNAs as a major epigenetic component of exceptional longevity in humans. By profiling the miRNAs in B-cells from Ashkenazi Jewish centenarians and 70-year-old controls without a longevity history, we found that the majority of differentially expressed miRNAs were upregulated in centenarians and predicted to modulate the IIS pathway. Notably, decreased IIS activity was found in B cells from centenarians who harbored these upregulated miRNAs. miR-142-3p, the top upregulated miRNA, was verified to dampen the IIS pathway by targeting multiple genes including *GNB2, AKT1S1, RHEB* and *FURIN*. Overexpression of miR-142-3p improved the stress resistance under genotoxicity and induced the impairment of cell cycle progression in IMR90 cells. Furthermore, mice injected with a miR-142-3p mimic showed reduced IIS signaling and improved longevity-associated phenotypes including enhanced stress resistance, improved diet/aging-induced glucose intolerance, and longevity-associated change of metabolic profile. These data suggest that miR-142-3p is involved in human longevity through regulating IIS-mediated pro-longevity effects. This study provides strong support for the use of miR-142-3p as a novel therapeutic to promote longevity or prevent aging/aging-related diseases in human.

## Introduction

Aging is a multifactorial process characterized by a progressive loss of physiological integrity, decline in tissue function and increased risk of multiple age-related pathologies^1^. Studies in model organisms have demonstrated that aging and longevity are influenced by genetic, epigenetic and environmental factors through multiple conserved signaling pathways across species^2^. The first established genetic pathway was the insulin/insulin-like growth factor-1 (IGF-1) signaling (IIS) pathway, identified through loss-of-function mutations in *daf-2* and *age-1* that reduces IIS signaling in *C. elegans*, leading to extended longevity^3, 4^. Mutations or interventions downregulating the activity of this pathway also were demonstrated to extend the lifespan in flies and mice^5, 6^. In humans, genetic variations in IGF1R are associated with and validated in exceptional longevity^7, 8^. Other genetic pathways, including rapamycin (mTOR), AMP-activating protein kinase (AMPK), and sirtuin signaling, also have been identified to modulate lifespan across species^1, 9^.

MicroRNAs (miRNAs) are important epigenetic regulators of biological mechanisms that are relevant to aging and longevity. miRNAs are small non-coding RNA species that post-transcriptionally regulate gene expression by inducing translational repression or mRNA degradation^10^. The miRNA regulatory network is extensive, as each miRNA can modulate multiple mRNA targets and each mRNA may be targeted by different miRNAs^11^. It is estimated that miRNAs participate in modulation of up to 60% protein-coding genes in mammals and have influence on a wide range of biological functions, such as stem cell self-renewal, cell proliferation, apoptosis, and metabolism^12–14^.

Multiple miRNAs such as lin-4, miR-34 and let-7 directly regulate life span of both of *C. elegans* and *Drosophila* positively and negatively^15–17^. Some of these longevity modulating miRNAs control the expression of genes involved in major conserved pathways that impact life span, such as the IIS^18^. Moreover, miRNAs were shown to be mediators of the longevity phenotype in Ames dwarf mice^19^, implicating miRNAs in mammalian longevity. Recently, transgenic expression of miR-17 was reported to extend lifespan and inhibits cellular senescence in mice. However, it is important to note that the role of miR-17 in promoting longevity is still controversial since it is generally considered to be an oncogene^20^. Since a significant number of miRNAs are evolutionarily conserved^21^, it is conceivable that miRNAs are also involved in the regulation of human longevity. Indeed, several miRNAs in humans (e.g., miR-29b and miR-143) are known to target the components of IGF-1 signaling and have been linked to human aging-related disorders such as diabetes and neurodegenerative disease^22, 23^. Recent studies further showed alterations in miRNAs expression profiles in long-lived individuals and identified multiple longevity-associated candidate miRNAs^24, 25^. However, the role(s) of miRNAs in human longevity and the underlying mechanism(s) is still unclear.

Here, to identify potential longevity-associated miRNAs, we performed a comprehensive miRNA transcriptome analysis in B-cells from centenarians, which are the unique, genetically homogenous, populations of Ashkenazi Jewish (AJ). As controls, 70 year old without a longevity history were used. We found that the far majority of differentially expressed miRNAs were significantly upregulated in centenarians as compared to controls. Many of these miRNAs are linked to regulation of the IIS signaling pathway. In particular, we demonstrate that miR-142-3p, one of the top upregulated miRNAs in centenarians, downregulated the IIS pathway through targeting multiple genes involved in IIS pathway both in cell culture and in mouse aging models. These data provide mechanistic insight into the regulation of the IIS pathway as a conserved aging pathway and reveal a novel role for miR-142-3p in human longevity.

## Results

### Differentially expressed miRNAs in centenarians are associated with IIS pathway

To discover the longevity-associated miRNAs, we performed a comprehensive miRNA transcriptome of B-cells established from 20 centenarians (mean age 101.8) and 20 elderly controls without a history of longevity (mean age 72.5). After differential analysis, there were a total of 49 miRNAs significantly differentially expressed between controls and centenarians (fold change > 2, False Discovery Rate < 0.05), and 44 of them showed increased expression in centenarians (**Fig. 1a, Supplementary Table 1**). Among these upregulated miRNAs, miR-142-5p and miR-142-3p showed the strongest significance and the highest difference between centenarians and controls, with the fold change of 43.37 and 32.30, respectively. The relative expression of these miRNAs in each control and centenarian subject is shown in **Fig. 1b**. Remarkably, the expression pattern of the 44 upregulated miRNAs clearly divided these centenarians into 2 groups, one (9/20) with a common enrichment signature of these longevity-associated miRNA, while the other group with no such signature. Consistently, PCA analysis also showed the centenarians with the signature were clearly separated from the control group and the centenarians without the signature (**Fig. 1c**). This expression pattern of the miRNAs was further confirmed by RT-qPCR analysis of 8 differentially expressed miRNAs (**Extended Data Fig. 1**). These results suggest that centenarians with the signature may share a common underlying mechanism related to their exceptional longevity, which can be attributed to some of those upregulated miRNAs. Interestingly, some of these centenarians-enriched miRNAs including miR-19b, miR-29b, miR-106b, and miR-142-5p, miR-142-3p, have been reported to be downregulated during human or murine aging^26–28^. To identify the possible pathways targeted by centenarian-enriched miRNAs, we used the DIANA-microT 3.0 target prediction program and found that P13K-Akt and FoxO signaling pathways were two of the top putative pathways targeted by these longevity-associated miRNAs (**Fig. 1d**). P13K-Akt and FoxO are part of the IIS signaling pathway, which is the most conserved aging-controlling pathway. To verity if the IIS signaling activity was changed in centenarians with upregulated miRNA signature, we compared the extent of AKT phosphorylation in B cells between centenarians with and without the signature. We found that centenarians who harbored the upregulated miRNA signature showed decreased IIS activity as compared to those without such signature (**Fig. 1e, f**). This data suggests that these miRNAs contribute to human longevity by regulating IIS activity.

**Figure 1.**
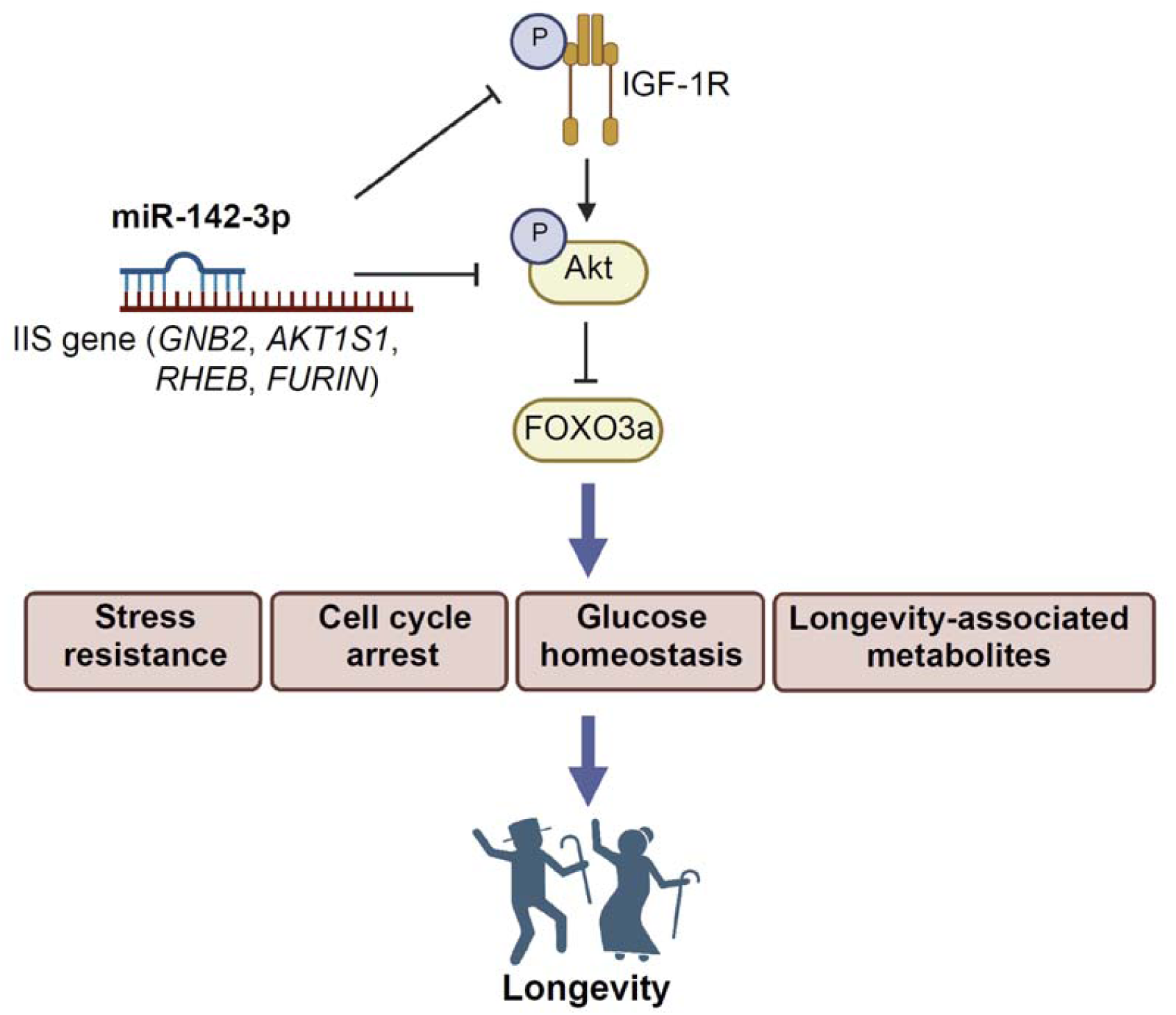
Differentially expressed miRNAs between controls with 70s and centenarians. (a) Volcano Plot depicting differentially expressed miRNAs between centenarians (n=20) and elderly controls (n=20) from small RNA-sequencing, discriminated based on fold change > 2, False Discovery Rate < 0.05. Colored dots correspond to miRNAs significanly enriched in centenarians (red dots), or significantly enriche in elderly controls (blue), or not significant in either groups (grey). (b) Heatmap showing relative expression of differentially expressed miRNAs (FDR < 0.05, Fold change >2) miRNAs in each control and centenarian subject. (c) Principal component analysis (PCA) score plot showing the seperarion of centenarians with and without signature based on all miRNAs detected from small RNA-sequencing. (d) Top putative pathways targeted by centenarian-enriched miRNAs based on DIANA-microT 3.0 target prediction program. (e) The protein level of phospho-AKT (Ser473) in the group harboring up-regulated miRNA signature within centenarians (f) The quantification analysis for the protein level of phospho-AKT (Ser473). Data are expressed as mean ± SEM. *p < 0.05; **p <0.01; ***p < 0.001.

### Reduced IIS activity by miR-142-3p in *in vitro* cell culture system

Among the centenarian-enriched miRNAs discovered in B cells, we found that 9 of them, including hsa-miR-142-3p, hsa-miR-101-3p and hsa-miR-301b. (**Supplementary Table 2**), also were significantly upregulated in plasma from centenarians, indicating they were longevity-associated circulating miRNAs that may have systemic effects. The ability of the top candidate miRNA, miR-142-3p to regulate IIS activity was examined by transfecting human breast cancer MCF7 cells with the miRNA mimic. Transfection of miR-142-3p significantly decreased levels of p-AKT and other components of the IIS pathway including p-IGF-1R and p-FOXO3A (**Fig. 2a, b**). Consistently, overexpression of miR-142-3p in human fibroblast IMR90 cell line also led to decreased p-AKT level (**Extended Data Fig. 2a, b**). These data suggest that miR-142-3p is a functional miRNA that could promote human longevity by dampening the IIS activity. Additionally, we compared the gene expression levels of miR-142-3p in offspring of AJ centenarians and age-matched controls in different age groups including 60s and 80s and found that miR-142-3p was significantly enriched in offspring of centenarians than in age-matched controls (**Extended Data Fig. 2c**), implicating that high expression of miR-142-3p as a possible heritable longevity-associated feature.

**Figure 2.**
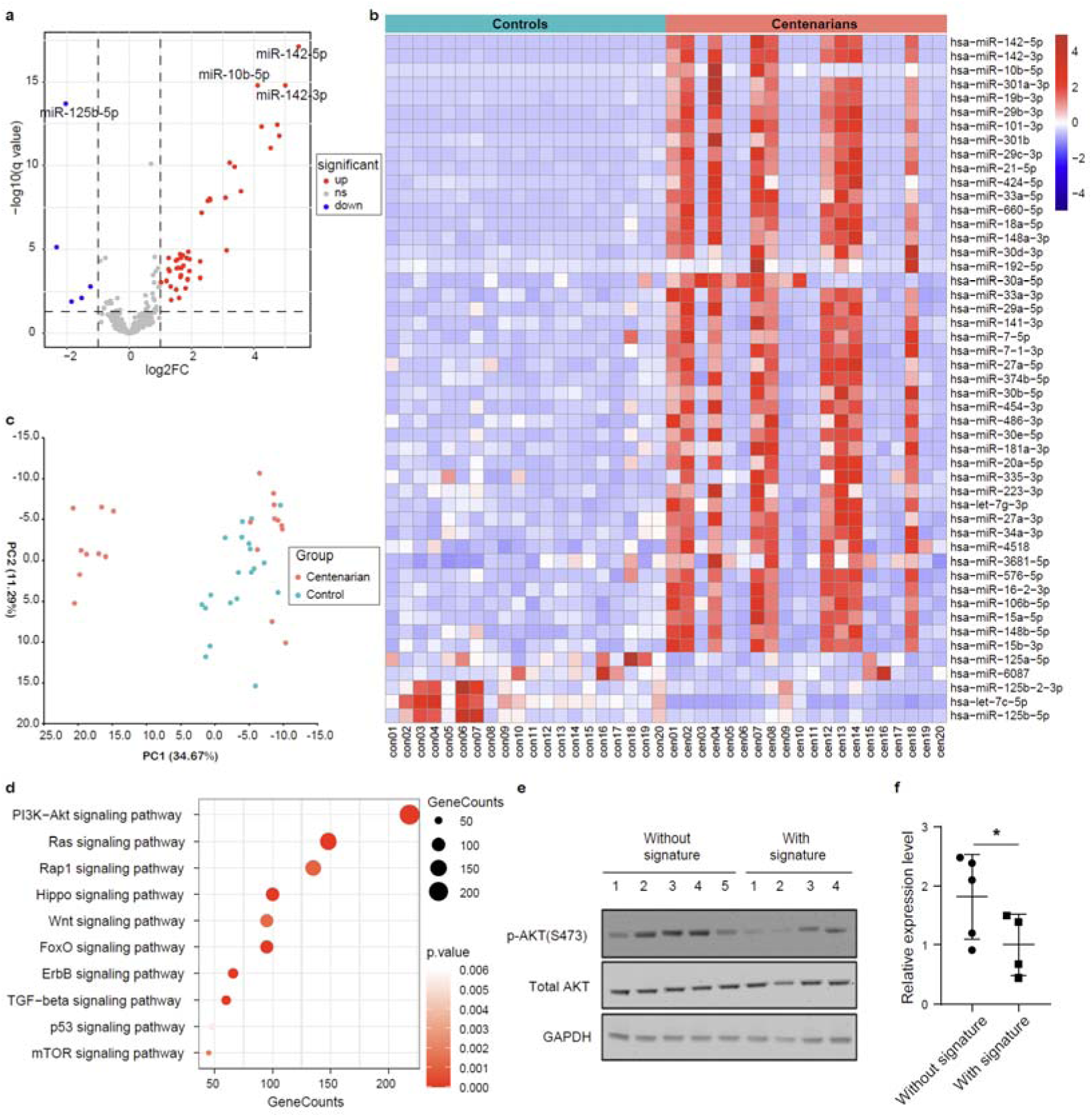
Reduced IIS activity and IIS pathway-involved genes regulated by longevity-associated miRNA miR-142-3p. (a) The protein level of IIS molecules after miR-142-3p transfection into MCF7 cells. (b) The quantification analysis for the protein level of IIS molecules from (a). (c-d) The GSEA enrichment plot for regulation of IIS (c) and DNA repair (d), showing the profile of the running ES Score and positions of gene set members on the rank-ordered list. (e) Heatmaps showing the relative expression level of gene targets of miR-142-3p in miR-142-3p transfected MCF7 cells and control cells based on RNA-seq data. (f) Enriched genes after pull-down assay in biotinylated miR-142-3p transfected MCF7 cells. (E) Summary of the IIS pathway and genes targeted by miR-142-3p directly or indirectly. Data are expressed as mean ± SEM. *p < 0.05; **p <0.01; ***p < 0.001.

### IIS pathway-involved genes targeted by miR-142-3p

To identify the targets of miR-142-3p to reduce IIS activity, we performed RNA-sequencing on miR-142-3p-transfected MCF7 cells (n=3) with cel-miR-67-transfected MCF7 cells used as a control (n=3) and identified the differentially expressed genes (DEGs, FDR < 0.05, Fold Change > 2, **Extended Data Fig. 2d**). Several aging-related pathways such as homology directed repair, chromosome maintenance and telomere maintenance were significantly altered (FDR < 0.05) in miR-142-3p overexpressed cells compared with controls (**Extended Data Fig. 2e**). Specifically, gene set enrichment analysis revealed that the IIS pathway was significantly downregulated while DNA repair pathways were significantly upregulated in miR-142-3p overexpressed cells (**Fig. 2c, d**). By overlapping the downregulated DEGs with miR-142-3p targets predicted by either Targetscan or DIANA algorithms, we identified 137 putative miR-142-3p targets **(Extended DataFig. 2f and Supplementary Table 3**). Among them were genes known to be involved in the IIS pathway including *GNB2*, *INPP5A*, *ITPR3*, *AKT1S1* and *RHEB* (**Fig. 2e)**. The gene expression levels of the 5 predicted targets were further confirmed by RT-qPCR, consistent with the RNA-seq data (**Extended Data Fig. 2g**).

To verify the direct binding of miR-142-3p to these predicted targeted genes, we performed a miR pull down assay in MCF7 cells transfected with biotinylated miR-142-3p (bi-miR-142-3p) or biotinylated cel-miR-67 (bi-cel-miR-67) as a control. Results showed that *GNB2*, *AKT1S1*, *INPP5A* and *ITPR3* were significantly enriched in streptavidin-pulled down samples from bi-miR-142-3p transfected cells, suggesting that miR-142-3p negatively regulates IIS pathway by directly targeting these four genes (**Fig. 2f**). However, *RHEB* were not enriched on bi-miR-142-3p, suggesting that *RHEB* is not a direct target of miR-142-3p (**Fig. 2f**). These data indicated that miR-142-3p can significantly reduce IIS signaling pathway by modulating multiple genes directly or indirectly (**Fig. 2g**).

Interestingly, there was no change in the level of the IGF-1R mRNA in miR-142-3p transfected MCF7 cells, but Western blotting showed IGF-1R protein level was significantly decreased, which also contributed to impaired IIS (**Fig. 2a, b**). It was known that pre-mature IGF-1R (pro-IGF-1R) can be cleaved by protein convertase *FURIN* to produce IGF-1R mature form (IGF-1R-beta-subunit)^29^. The level of FURIN protein was reduced in miR-142-3p transfected MCF7 cells (**Extended Data Fig. 2h**), which could explain the increased protein level of pre-mature IGF-1R and decreased production of mature IGF-1R. Based on the sequences at 3’UTR, FURIN was not predicted as the direct target of miR-142-3p. However, miR-142-3p dramatically repressed FURIN mRNA level in MCF7 cells, suggesting miR-142-3p indirectly regulate the transcriptional activity of FURIN and may have a broader impact on the regulation of other IIS genes.

### Beneficial effects of miR-142-3p on human normal fibroblast IMR90 cells

Impaired IIS extend lifespan through the modulation of numerous cellular processes in model organisms, including increased stress resistance and reduced cell cycle progression^30^. Therefore, to investigate if miR-142-3p can induce similar changes, we firstly measured stress resistance in human diploid fibroblast IMR90 cells transfected with miR-142-3p or cel-miR-67 as a control, under the exposure of different kinds of genotoxic stressors. After diquat treatment, IMR90 cells transfected with a miR-142-3p mimic showed a significantly higher cell viability and lower level of ROS when compared to the control group (p<0.05) (**Fig. 3a, b**). In addition, miR-142-3p transfection significantly decreased cell death as compared to control group following exposure to cadmium or etoposide (*p* <0.05) (**Extended Data Fig. 3**). These results indicate that miR-142-3p improves the stress resistance under genotoxic stress *in vitro*. Next, the impact of miR-142-3p on cell cycle progression was examined. The results showed transfection of IMR90 cells with the miR-142-3p mimic inhibited the transition from G0/G1 phase to S phase (*p* <0.05) (**Fig. 3c, d**). These results suggest that miR-142-3p impairs cell cycle progression, which is a similar result with our previous finding that IGF1R mutations in Igf1r-knock out mouse embryonic fibroblasts caused a delay in cell cycle progression^8^.

**Figure 3.**
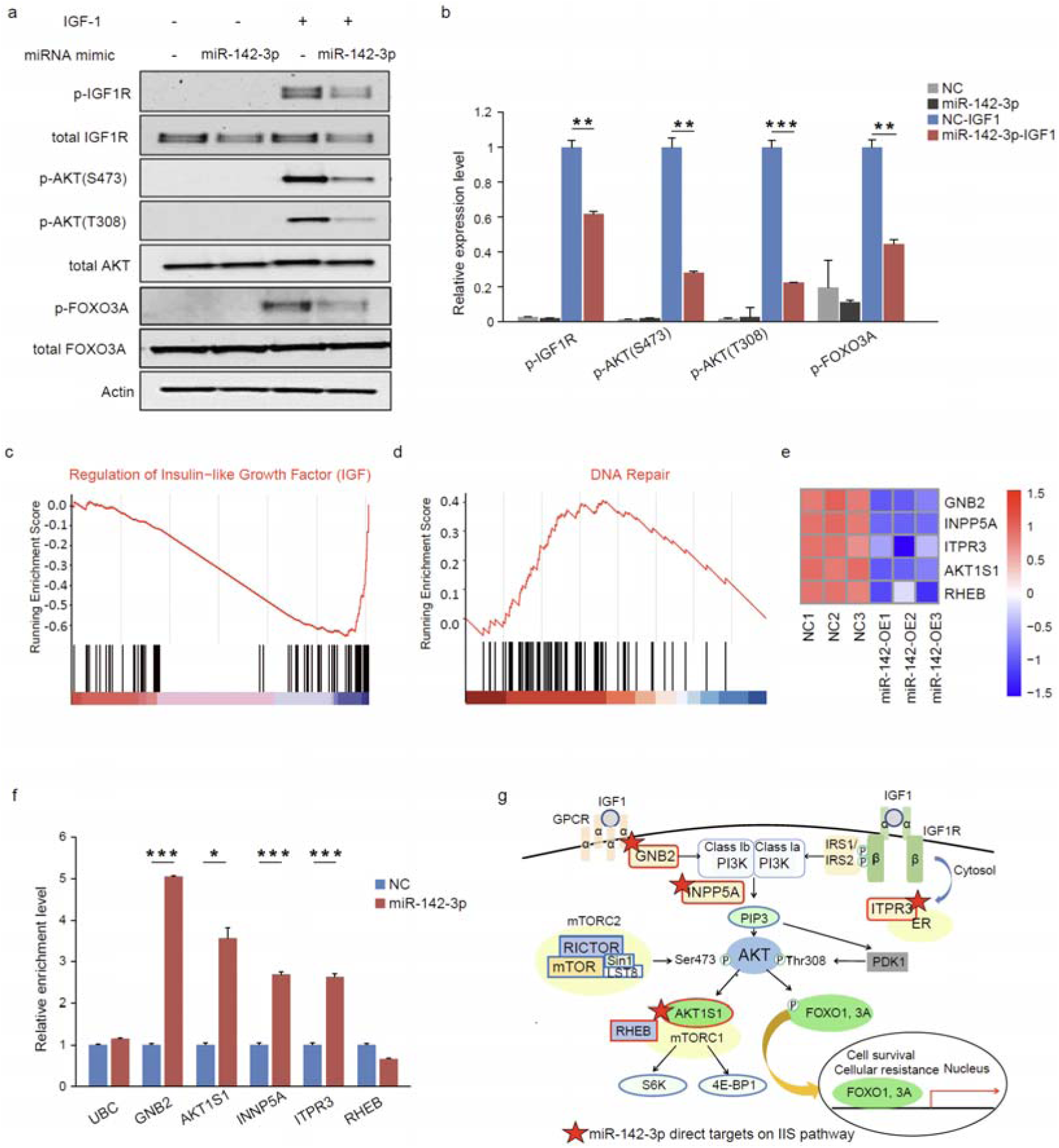
Beneficial effects of miR-142-3p on human normal fibroblast IMR90 cells. (a) Cell viability under different dose of diquat treatment in IMR90 cells transfected by miR-142-3p or cel-miR-67 using MTS assay. (b) Level of ROS after different dose of diquat treatment in IMR90 cells transfected by miR-142-3p or cel-miR-67 using FACS analysis. (c) Cell cycle analysis of IMR90 cells transfected by miR-142-3p or cel-miR-67 using FACS. The percentage of cells in G0/G1, S and G2/M phase is indicated by propidium iodide (PI) staining in combination with 5-ethynyl-2 deoxyuridine (EdU) incorporation. (d) Quantitative results of percentage of cells in G0/G1, S and G2/M form (c). Data are expressed as mean ± SEM. *p < 0.05; **p <0.01; ***p < 0.001.

### Reduced IIS activity in liver tissue of mice overexpressing miR-142-3p

To elucidate the role of miR-142-3p *in vivo*, we investigated the effect of this miRNA on IIS signaling pathway by intravenous injection of miR-142-3p mimics or cel-miR-67 as a control into mice daily for 6 days (**Fig. 4a**). Initially, the levels of miR-142-3p in serum and in tissues including liver, spleen muscle, heart, kidney, adipose and brain were measured. The levels of miR-142-3p were significantly increased in serum, liver, spleen and adipose of mice injected with miR-142-3p as compared to controls (**Extended Data Fig. 4**) with the greatest increase in liver (**Fig. 4b**). The effect on the IIS pathway was examined in liver tissue by measuring the levels of p-Igf1r and p-Akt, which were significantly reduced in mice injected with miR-142-3p as compared to controls (**Fig 4c, d**). In addition, the protein levels of Igf1r and Furin was significantly decreased in the liver tissue of miR-142-3p injected mice (**Fig 4c, d**), which is consistent to the results in MCF7 cells **(Extended Data Fig. 2h)**. Moreover, we found that the expression levels of target genes of miR-142-3p found *in vitro* including *GNB2*, *AKT1S1*, and *RHEB* were also significantly decreased in liver tissue of miR-142-3p-injected mice as compared to controls (**Fig. 4e**). Taken together, these data suggest that miR-142-3p could downregulate IIS activity by negatively regulating multiple molecules involved in the IIS pathway in mice.

**Figure 4.**
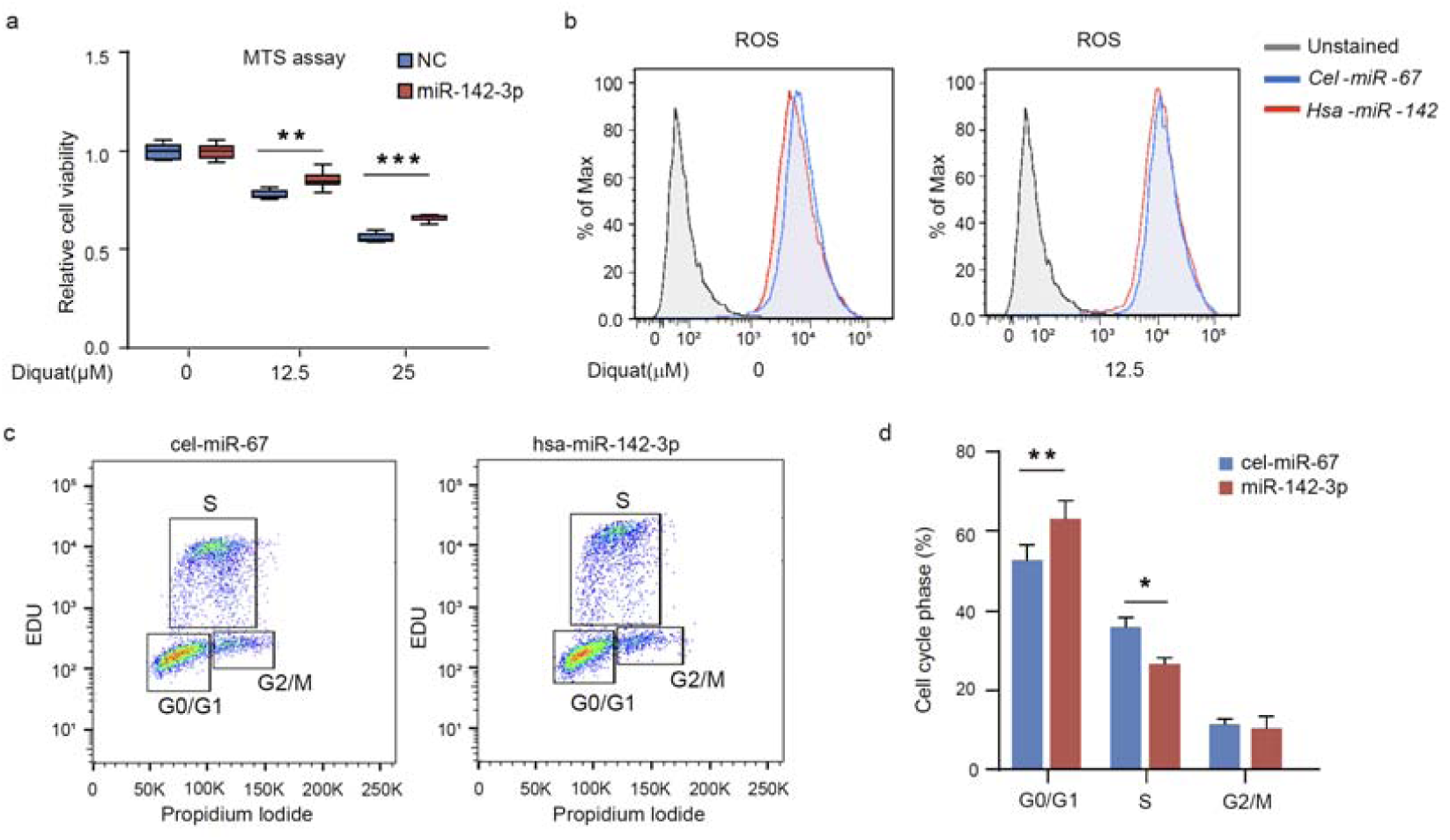
Reduced IIS activity by miR-142-3p overexpression in mice liver tissues. (a) Schematic model for miR-142-3p overexpression in mice by tail vein injection. (b) Relative expression level of miR-142-3p in liver tissue after injection for 6 days in miR-142-3p-overexpressing mice and control mice. (c) The protein level of IIS molecules in liver tissue after miR-142-3p injection for 6 days in miR-142-3p-overexpressing mice and control mice. (d) The quantification analysis of protein level of IIS molecules from (c). (e) The mRNA level of genes involved in IIS pathway in liver tissue of miR-142-3p injected mice. Data are expressed as mean ± SEM. *p < 0.05; **p <0.01; ***p < 0.001.

### Protective effect of miR-142-3p against genotoxic stress in *in vivo* mouse study

To see if miR-142-3p has a protective effect against genotoxic stressors *in vivo* miR-142-3p and control cel-miR-67 injected mice were exposed to diquat to induce oxidative stress. Analysis of tissue pathology by H&E staining revealed decreased liver extramedullary hematopoiesis (EMH), kidney vacuolation and spleen EMH, which are the indicators of toxicity in the pathological aspect, specifically in the miR-142-3p treated mice (**Fig. 5a-c**). Several detoxification-related enzyme classes such as cytochrome P450 (CYPs), UDP-glucuronosyltransferase (UGTs), and glutathione S-transferase (GSTs) are regulated by IIS and act in concert to dispose of toxic endobiotic or xenobiotic compounds (e.g., toxins, drugs, and carcinogens). Analysis of the expression levels of detoxification genes in liver using RT-qPCR demonstrated that *Cyp3a25, Cyp3a59, Gsta3, Gstp1, Gstm1, Gstm3* were upregulated by miR-142-3p injection (**Fig. 5d**). These data indicated that miR-142-3p has a protective role *in vivo* against genotoxic stressors through elevating expression of genes involved in detoxification.

**Figure 5.**
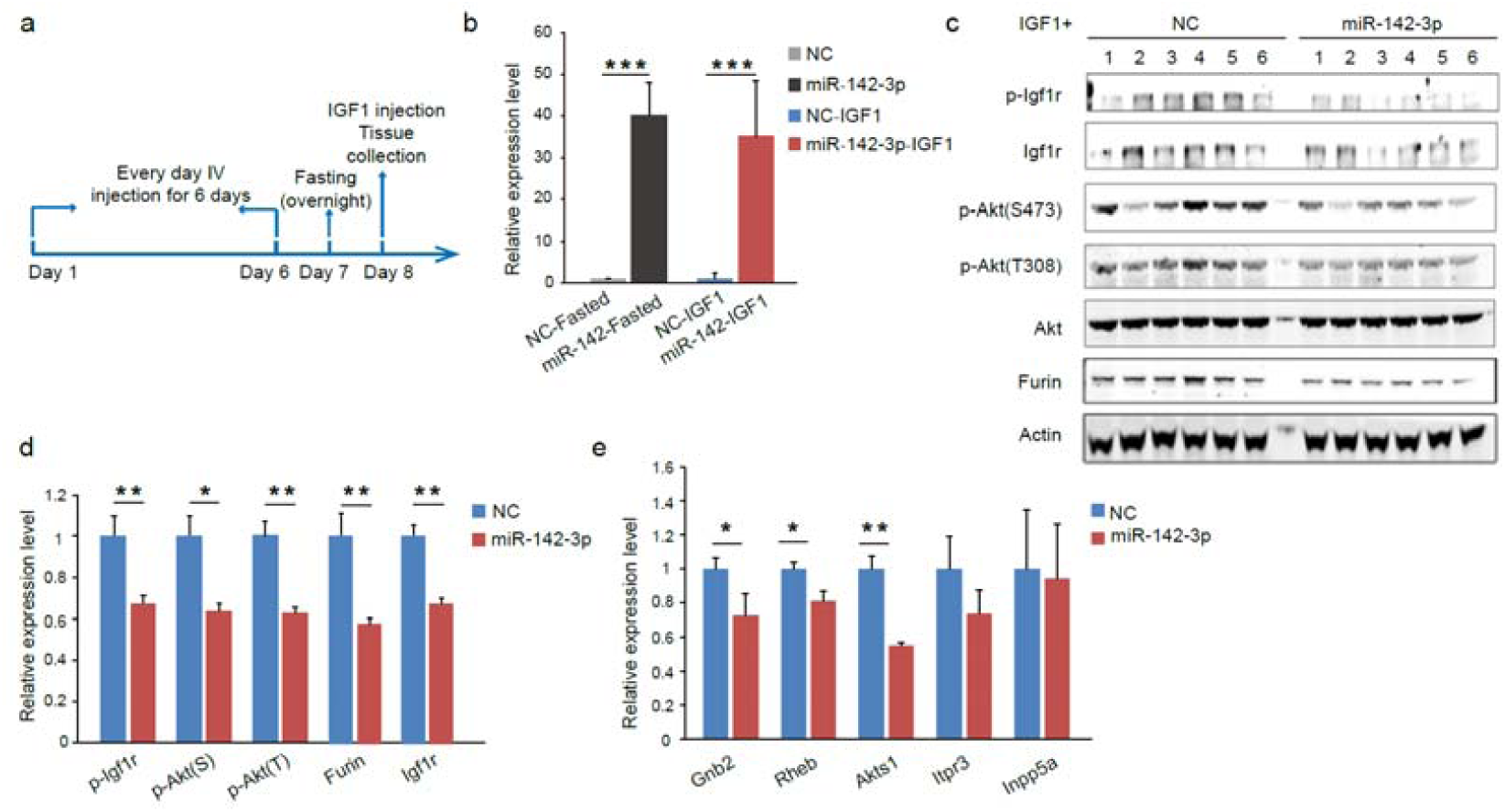
Improvement of stress resiswstance and glucose intolerance and liver metabolic alterations in miR-142-3p overexpressing mice. (a-c) H&E-staining and quantification results showing liver extramedullary hematopoiesis (EMH)(a), kidney vacuolation (b) and spleen EMH (c) in negative control-injected mice and miR-142-3p-injected mice after diquat treatment for 4 hours (n=5). (d) The mRNA level of genes involved in detoxification in liver tissue of miR-142-3p injected mice. (e-f) Oral glucose tolerance test (OGTT) (e) and intraperitoneal insulin tolerance test (ipITT) (f) in miR-142-3p-injected mice (n=9) and negative control-injected mice (n=10) under HFD for 4 weeks before the injection. (g-h) OGTT (J) and ipITT (K) in 5 months old mice injected by miR-142-3p (n=7) and negative control (n=8). (i-j) OGTT (i) and ipITT (j) in 18 months old mice injected by miR-142-3p (n=10) and negative control (n=11). Data are expressed as mean ± SEM. *p < 0.05; **p <0.01; ***p < 0.001. (k-l) OPLS-DA score plot of GC-MS (k) and LC-MS (l) in mouse liver. (m-n) Heatmap showing differently expressed metabolites from GC-MS (k) and LC-MS (l) analysis (VIP > 1, p < 0.05)

### Improvement of glucose homeostasis by miR-142-3p

Lifespan extending interventions targeting IIS signaling such as caloric restriction improve insulin sensitivity and glucose homeostasis in both human and mouse studies^31, 32^. To test whether the reduction of IIS activity by miR-142-3p in mice on a high-fat diet affects glucose homeostasis, we performed an oral glucose tolerance test (OGTT) and an intraperitoneal injection insulin tolerance test (ipITT) in mice injected with miR-142-3p or with cel-miR-67. Under a high fat diet, the mice treated with miR-142-3p showed significant lower glucose levels in blood at 60 min and 120 min as compared to control mice (**Fig. 5e**), although there was no difference on insulin tolerance from ipITT (**Fig. 5f**). These data suggest that miR-142-3p could improve diet-induced glucose intolerance. In addition, miR-142-3p-injected mice showed significantly lower (p=0.05) weight gain (%) than control mice at 3 and 4 weeks after the injection (**Extended Data Fig. 5a**). In addition, the respiratory exchange ratio in miR-142-3p-injected mice was significantly lower than the control mice during both light and dark phases (**Extended Data Fig. 5b**), suggesting that they tend to oxidize fat rather than carbohydrates for energy utilization. However, there is no significant difference in average day and night energy expenditure value or physical activity levels between these two groups (**Extended Data Fig. 5c-e**). These data suggest that miR-142-3p regulates glucose homeostasis and systemic energy homeostasis in mice on a high fat diet.

We also measured both glucose and insulin tolerance in young (5 months old) and old (18 months old) male mice after the injection with miR-142-3p or cel-miR-67. In young mice, there is no significant difference in glucose tolerance and insulin sensitivity between miR-142-3p-injected mice and control mice (**Fig. 5g, h**). In contrast, old mice injected with miR-142-3p showed significantly decreased blood glucose levels at 60- and 120-min post-glucose injection (**Fig. 5i**), suggesting improved glucose tolerance by miR-142-3p injection. Moreover, old mice injected with miR-142-3p also showed significant lower blood glucose level at 15-, 30- and 60-min post-insulin injection (**Fig. 5j**), suggesting miR-142-3p can increase insulin sensitivity with aging. Thus, these data indicate that miR-142-3p not only improves diet-induced glucose intolerance and but also has a beneficial effect on glucose homeostasis during aging.

### Alteration of metabolic profiles in liver tissue of mice overexpressing miR-142-3p

Previous studies have shown unique metabolic profiles in long-lived models including *C. elegans*^33, 34^ and long-lived human populations^35, 36^. Therefore, to see if there is any correlation in longevity metabolic profiles and miR-142-3p-overexpressing mice, GC– MS and LC-MS based metabolomics were performed in liver tissue of mice injected with miR-142-3p and cel-miR-67. According to the score plot of orthogonal projections to latent structures discriminant analysis (OPLS-DA), we found that the metabolic pattern of miR-142-3p-injected mice were separated from those of control mice based on both LC-MS and GC–MS matrix with satisfactory goodness of fit (**Fig. 5k, l and Extended Data Fig. 5f, g)**. To further refine the significantly different metabolites, standards of VIP >1 in supervised OPLS-DA model and *p* value <0.05 were used. From GC-MS data, 16 annotated small polar metabolites were significantly different in miR-142-3p-injected mice. Of them, 10 metabolites were enriched in miR-142-3p-injected mice, including multiple kinds of amino acids, such as alanine, lysine, threonine, and isoleucine (**Fig. 5m and Supplementary Table 4**), which have been reported to be down-regulated with age and involved in extending life span of *C. elegans*^33, 34^. From LC-MS analysis, 10 lipid metabolites including 8 glycerolphospholipids (PCs) and 2 sphingolipids (SMs) showed significant difference between miR-142-3p-injected and control mice (**Fig. 5n and Supplementary Table 5**). Of them, PC ae C34:2, PC ae C36:3, and PC ae C36:2 were up-regulated in the liver of miR-142-3p-overexpressing mice. Interestingly, these PCs were also reported to be enriched in blood of centenarians and offspring of nonagenarians when compared with controls in previous studies^35, 36^. Taken together, these results suggest that the alteration of metabolites by miR-142-3p could be involved in a pro-longevity mechanism.

## Discussion

Although miRNAs have been linked to healthspan and lifespan in models of aging, a possible role for miRNAs in human longevity has not been elucidated. Here we used a cohort of Ashkenazi Jewish (AJ) centenarians, compared to elder controls at 70s without longevity history, to identify a subset of miRNAs differentially expressed in B cells and circulating in centenarians. We further demonstrate that miR-142-3p plays a key role in modulating the IIS signaling pathway *in vitro* and *in vivo* that potentially regulates longevity.

Interestingly, most of the miRNAs that are differentially expressed in B cells between centenarians and controls were upregulated, not downregulated in centenarians. This result is consistent with Serna et al.’ findings, which observed that centenarians but not octogenarians upregulate the expression of miRNAs in peripheral blood mononuclear cells (PBMCs)^24^. This observation suggests that it is an increase and not a decrease in miRNA function that is important for longevity. Importantly, the upregulation of miRNAs in long-lived individuals is consistent with the observation that DROSHA, DICER, and other factors involved in the control of these miRNAs biogenesis genes are upregulated in centenarians when compared with octogenarians^37^. Interestingly, we found that these miRNAs were overexpressed only in a subset of the centenarians, referred to as centenarians with signature. Also, IIS activity was significantly decreased in these centenarians with signature, consistent with the possibility that these miRNAs regulate human longevity by modulating the IIS signaling pathway. The IIS signaling pathway is the best characterized longevity-associated pathway which has been known to affect health span from yeasts to mammals^38^. Several variants in genes within the insulin/IGF-1-signaling pathway, such as FOXO3A and AKT1, have been discovered to be robustly associated with human longevity^39^. In particular, our previous study demonstrated that functional mutations in the IGF-1R gene are overrepresented in a cohort of Ashkenazi Jewish (AJ) centenarians, resulting in an altered IIS signaling pathway^40^. Moreover, it was verified that the longevity-associated IGF-1R mutations attenuated the IGF1 signaling and caused a delay in cell cycle progression in Igf1r knock-out mouse embryonic fibroblasts (MEFs)^8^. This present study has extended the regulation of IIS signaling from variants in specific proteins to miRNAs that are upregulated specifically in centenarians.

Of the differentially regulated miRNAs, miR-142-3p was found to be the top upregulated miRNAs in B cells and blood from centenarians. miR-142-3p serves as a guide strand of pre-mature miR-142, is evolutionally conserved in vertebrates and is known to be highly expressed in hematopoietic cells^41^ where it regulates cell fate decisions^42, 43^. In addition, miR-142-3p has emerged as a multifaceted regulator in development and diseases^44^ and has been reported to balance self-renewal and differentiation in mESCs via regulating KRAS/ERK signaling^45^. Recently, downregulation of miR-142-3p expression was found in peritoneal macrophages in aged mice where it may contribute to IL-6-associated aging disorders^28^. Moreover, the age-dependent decline of miR-142-3p levels was observed in a human association study in 1334 healthy individuals ranging in age from 30 to 90 years^46^. Interestingly, we found that miR-142-3p was significantly upregulated not only in centenarians, but also in their offspring as compared to age-matched controls. Offspring of centenarians always have a markedly reduced prevalence of age-related diseases as compared to unrelated age-matched controls^47^ and show centenarian-enriched genotypes and molecular phenotypes, indicating the heritability of exceptional longevity^48, 49^. For example, lower circulating IGF-I levels were found in both centenarians and their offspring, shown to be associated with human longevity^50, 51^. Thus, our results suggest that miR-142-3p is a longevity-associated miRNA which decreases with aging but remarkably increases in centenarians.

We verified the downregulation the IIS signaling pathway by miR-142-3p both *in vitro* and *in vivo*. Along miR-142-3p dampens IIS activity by targeting multiple genes involved in IIS pathway, miR-142-3p also decreased the IGF-1R processing by decreasing the expression level of *FURIN*, important for processing of IGF-1R. Inhibition of *FURIN* reduces the incidence, size and vascularization of tumor development in transplanted mice^52^. Recently, IGF-1R monoclonal antibodies have been also proved to induce delayed aging, showing the consistency with genetic models of IGF-1R heterozygosity^53^. Therefore, our data indicate that by targeting multiple IIS related genes and inhibiting IGF-1R processing, modulation of miR-142-3p might be a novel and efficient approach to combat aging/aging-related diseases or promote longevity.

We also demonstrated beneficial effects of miR-142-3p on the response to genotoxic stress both in cell culture and *in vivo*. A decrease in the IIS pathway has been observed to lead to an improved response to cellular stress and an increased lifespan in several animal species including worms, fruit flies and mice^30, 54^. Our data showed that stress resistance was enhanced by miR-142-3p in normal human fibroblasts IMR90 against genotoxic agents such as diquat and cadmium. This result is similar to skin-derived fibroblasts from long-lived dwarf mice that are resistant to multiple forms of cellular stress^55^. Moreover, analysis of pathology of H&E-stained mouse tissues showed that non-neoplastic lesions such as liver EMH, spleen EMH and kidney vacuolation which are typically associated with pathologic condition and most often seen in animals treated with a toxicant were decreased by miR-142-3p. This suggests that miR-142-3p has a protective role against a toxic environment, which also could be important for human longevity. Several detoxification related gene classes appear to be regulated by the IIS pathway. For example, upregulation of genes encoding the cytochrome P450 (CYPs), short-chain dehydrogenase/reductase (SDRs), and UDP-glucuronosyltransferase (UGTs) has been observed in *daf-2* adults and dauer larvae^56^. Moreover, there is upregulation of detoxification related gene group, the glutathione S-transferases (GSTs), in long-lived mouse models such as little mice^57^. In this study, we found that some detoxification genes including *CYP3a25*, *GSTA3*, *GSTM1* and *GSTM3*, which are known to be upregulated in long-lived little mice, were upregulated by miR-142-3p overexpression in liver tissue. These results suggest that miR-142-3p could involve in longevity through the activation of IIS-regulated detoxification genes against the toxicant exposure.

In addition, we demonstrated that miR-142-3p can regulate glucose homeostasis in mice on a high-fat diet and in aged mice. With obesity or age, insulin sensitivity progressively declines, which significantly contributes to the increased incidence of type 2 diabetes mellitus in older people^58, 59^. Remarkably, centenarians and their offspring were found to exhibit preserved glucose tolerance and insulin sensitivity, leading to a reduced risk of diabetes^47, 60^. In addition, offspring of long-lived nonagenarian siblings also were found to have enhanced insulin sensitivity and better glucose tolerance compared to a control group of similar age and body composition^61^. This suggests that people with exceptionally long lifespan may have higher insulin sensitivity throughout their lifespan with slower age-related decline of insulin activity. Moreover, many growth hormone (GH)-deficient or GH-insensitive human and mouse populations appear to be protected from diabetes mellitus^30, 62^. Here, our data showed that diet- and aging-induced glucose intolerance were both improved in mice treated with miR-142-3p. This effect of miR-142-3p overexpression was similar to caloric restriction, which is a well-known lifespan extension intervention inhibiting IGF-I-dependent signaling pathway. Caloric restriction was found to reduces plasma levels of IGF-I and increase insulin sensitivity in both rodents^31^ and human^32^. Thus, miR-142-3p may have a role of pro-longevity to protect from metabolic diseases under diet or aging by downregulation of IIS pathway.

In summary, our study implicates miR-142-3p as a potential longevity-associated miRNA targeting genes encoding proteins in the IIS pathway and improving glucose homeostasis, stress resistance and longevity-associated metabolites (**Fig. 6**). The identification of longevity-associated miRNAs in centenarians and their multiple mRNAs targets makes these molecules interesting therapeutic candidates to treat aging and age-related disorders. Importantly, advances in technologies to deliver RNA molecules *in vivo* have made miRNA-based therapeutics feasible. Therefore, miR-142-3p should be considered as a potential candidate to modulate IIS pathway, a conserved aging pathway in human.

**Figure 6.**
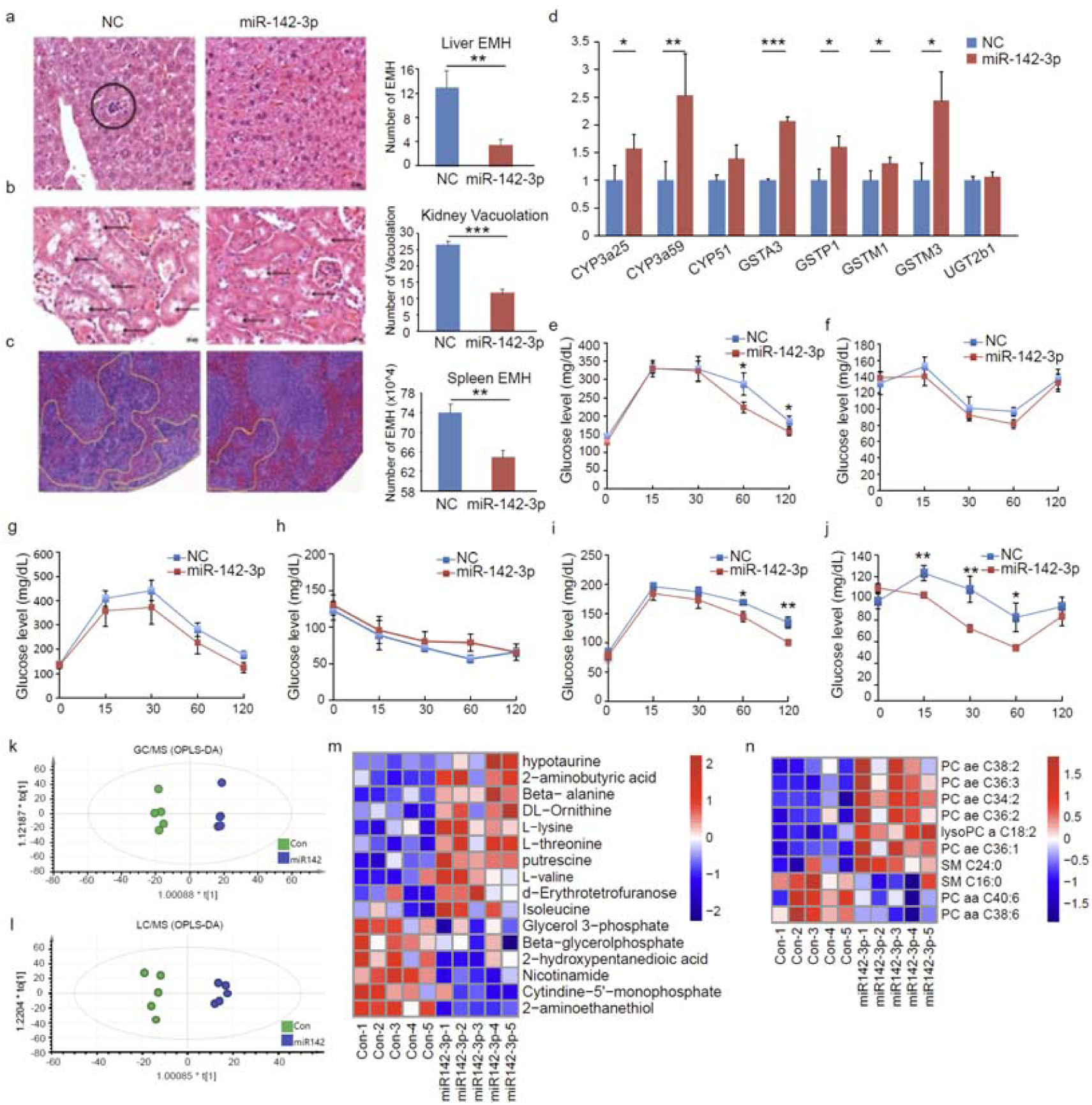
Proposed mechanism of miR-142-3p to promote longevity.

## Methods

### Population and sample collection

All individuals are enrolled in the Longevity Genes Project, and were recruited as described previously^63^. Informed written consent was obtained in accordance with the policy of the Committee on Clinical Investigations of the Albert Einstein College of Medicine. All blood samples were processed at the General Clinical Research Center at the Albert Einstein College of Medicine to produce EBV transformed immortalized B-cells as a source of RNA. Total RNA was extracted from immortalized B-cell lines established from centenarians and controls using TRIZOL reagent as recommended by the manufacturer (Invitrogen).

### Animals and experimental design

All procedures involving animals were approved by the Institutional Animal Care and Use Committee (IACUC) of Albert Einstein College of Medicine. All mice used in this study were male C57BL/6 and determined to be tumor-free and lacked other obvious signs of macropathology at the time of sacrifice. The C57BL/6 mice were purchased from Charles River Laboratories and kept at 23 °C in a humidified atmosphere with food and water ad libitum under standard 12-hour light-dark condition.

To study the effect of miR-142-3p on insulin/IGF-1 signaling pathway *in vivo*, PEI-complexed miR-142-3p or PEI-complexed cel-miR-67 was intravenously injected to 5-month-old mice every day through tail vein for 6 days. After fasting overnight, mice were given an intraperitoneal injection of 1 mg/kg body weight of rhIGF-1 (Austral Biologicals, San Ramon, CA, USA) or an equivalent volume of sterile saline. Ten min post injection, serum and tissues including liver, spleen, muscle et al. was collected and frozen in liquid nitrogen. Liver tissue homogenates were prepared, and protein concentration was determined by BCA assay.

To study the effect of miR-142-3p against genotoxic stress *in vivo*, 5-month-old C57BL/6 mice were used. 6 days after intravenous injection of PEI-complexed miR-142-3p (n=5) or PEI-complexed cel-miR-67 as control (n=5), 50 mg/kg diquat was exposed to mice by intraperitoneal injection to induce oxidative stress. Four hours after the injection, mice were sacrificed to collect the liver, kidney, and spleen tissues.

To see the impact of miR-142-3p on glucose homeostasis under high-fat diet, 5-month-old C57BL/6 mice were fed a 60 % HFD (D12492, Research Diets) starting at 8 weeks of age and continuing for 4 weeks. Then, PEI-complexed miR-142-3p and PEI-complexed cel-miR-67 were administered by intravenous injection for 6 days through the tail vein. To see the impact of miR-142-3p on glucose homeostasis under aging, young (5 months old) and old (18 months old) male were injected with PEI-complexed miR-142-3p and PEI-complexed cel-miR-67 as controls for 6 days.

### miRNAs, cell lines and cell culture

Chemically synthesized miRNAs mimics were purchased from GE healthcare Dharmacon. All cell lines were obtained from the American type culture collection (ATCC) and authenticated by the vendor. Cells were cultivated in a humidified incubator under standard conditions (37 °C, 5 % CO_2_) in RPMI or EMEM supplemented with 10 % fetal bovin serum.

### Ethics

Experimental research reported in this manuscript has been performed with the approval of the Committee on Clinical Investigations of the Albert Einstein College of Medicine. Research carried out in this manuscript is in compliance with the Helsinki Declaration (http://www.wma.net/e/policy/b3.htm).

### MiRNA-sequencing

2 μg of total RNA from each tissue sample and 10 ng of total RNA from each serum sample were used for small RNA cDNA library preparation. Using Illumina TruSeq Small RNA sample preparation kit, barcoded 3’-adapters and 5’-adapters were ligated to RNA in each sample. Synthesized cDNA amplified up to 15 cycles using indexes for multiplexing. Amplified DNAs were selected by the excision (145-160 bp) on 10 % TBE polyacrylamide gel and purified. Eluted library was qualified using bioanalyzer and quantified using Qubit. Sequencing was performed on an Illumina Hiseq 2000 analyzer.

### RNA library preparation and sequencing

Total RNA was isolated with the RNeasy Mini Plus Kit (Qiagen) and the quality of RNA (RIN>9) was checked using Bioanalyzer (Agilent). Strand specific RNA sequencing library were prepared by BGI using MGI rRNA removal kits & MGI directional kits, and were further sequenced on MGI’s with PE100 on the MGISEQ-2000 platform.

### Data analysis of miRNA sequencing and mRNA sequencing

As described previously, the sequencing data, in FASTQ format, were trimmed of adapter sequences using QUART. Sequences from each of the samples were aligned to the known human miRNAs present in hg19 using version 20 of mirBase. Analysis of the resulting read counts was performed in EdgeR. Read counts were normalized by library size, giving values in reads per million (rpm), as described previously. After normalization, any miRNAs with fewer than 10 rpm in more than 33 % of the samples were excluded. For RNA-sequencing, the reads were aligned with STAR (version 2.7.0), and genes annotated in GENCODE (release 38) were quantified with featureCounts in Subread package (version 2.0.3). Normalization and differential expression were performed using DESeq2 (version 1.32.0). Gene set enrichment analysis were assessed by clusterProfiler 4.0 and GSEA software (Broad Institute).

### Western blot

Total cell protein (15–20 μg) was used for Western blotting. Samples were resolved in 3-8 % NuPage Norvex tris-acetate Protein Gels (Life Technologies) and transferred to PVDF membranes (Life Technologies). The membranes were immersed in 0.5 % skim milk in TBS containing 0.1 % Tween 20 for 2 hours and probed overnight at 4 °C with primary polyclonal and monoclonal antibody against p-IGF1R (#3024), total IGF1R (#3027), p-AKT (S473, #4060), p-AKT (T308, #4056), total AKT (#4691), p-FOXO3a (#9466), total FOXO3a (#2497) from Cell Signaling Technology, FURIN (sc-133142) from Santa Cruz and beta actin (Ab8227) from Abcam. Blots were washed in TBS containing 0.1 % Tween20 and labeled with horseradish peroxidase (HRP)-conjugated secondary anti-mouse or anti-rabbit antibody (Cell Signaling Technology, Beverly, MA). Proteins were enhanced by chemiluminescence (Amersham ECL plus Western Blotting detection system, Fairfield, CT) for visualization. The protein expression levels were expressed relative to beta-actin levels.

### Quantitative RT-PCR analysis

Total RNA was isolated from B-cells or mice tissue using RNA isolation kit (Qiagen, Valencia, CA) and then converted to complementary DNA using TaqMan Reverse Transcription kit (Applied Biosystems, Foster City, CA) with microRNA specific RT primer (Applied Biosystems). A TaqMan^®^ microRNA assay was performed using AB StepOne^TM^ real-time PCR system to quantify relative miRNAs expression in these samples. The 20 µl total volume final reaction mixture consisted of 1 µl of TaqMan microRNA specific primer, 10 µl of 2x Universal Master Mix with no AmpErase^®^ UNG (Applied Biosystems) and 1.3 µl of complementary DNA. PCR was performed using the following conditions: 50 °C for 2 min, 95 °C for 10 min, 40 cycles of 95 °C for 15 sec, and 60 °C for 1 min. All reactions were run in duplicate to reduce confounding variance. U6 snRNA (Applied Biosystems) was used as an internal control. Means from different conditions were compared using the Student’s t-test. A significance threshold of *p* < 0.05 was used.

### Pull-down assay

MCF7 cells were transfected with Bi-miR-142-3p or Bi-cel-miR-67 (Dharmacon) in six well plates. Twenty-four hours later, the cells from wells were pelleted at 5000g. After washing twice with PBS, cell pellets were resuspended in 0.7 ml lysis buffer (20 mM Tris (pH 7.5), 100 mM KCl, 5 mM MgCl2, 0.3 % NP-40, 50 U of RNase OUT (Invitrogen), complete miniprotease inhibitor cocktail (Roche Applied Science)), and incubated on ice for 5 min. The cytoplasmic lysate was isolated by centrifugation at 10,000 xg for 10 min. Streptavidin-coated magnetic beads (Invitrogen) were blocked for 2 hr at 4 °C in lysis buffer containing 1 mg/ml yeast tRNA (Ambion) and washed twice with 1 ml lysis buffer. Cytoplasmic lysate was added to the beads and incubated overnight at 4 °C before the beads were washed five times with 1 ml lysis buffer. RNA bound to the beads (pull-down RNA) or from 10% of the extract (input RNA), was isolated using Trizol LS reagent (Invitrogen). The level of mRNA in the bi-miR-142-3p or bi-cel-miR-67 control pull-down was quantified by qRT-PCR. For qRT-PCR, mRNA levels were normalized to a housekeeping gene, *GAPDH*. The enrichment ratio of the control-normalized pull-down RNA to the control-normalized input levels was then calculated.

### Cell cycle

Cells harvested with trypsin were washed with phosphate buffered saline (PBS) and fixed by dropping them into 300 µl of 70 % ethanol pre-chilled to −20 °C. Cells were incubated in -4 °C for at least 1 hour. After washing with PBS, cells were treated with 10 mg/ml RNaseA (final concentration being 0.2-0.5 mg/ml) and incubated in 37 °C for 1 hour. To stain the cells for DNA content, 1 mg/ml propidium iodide (PI) along (final concentration being 10 µg/ml) was added to the cell solution. Cells were then analyzed by flow cytometry using CantoII software. Cylchred software was used in analyzing the percentage of cells in each phase of the cell cycle. Ten thousand cells were analyzed for each sample.

### Cell viability

IMR90 cells were transfected with 50 nM of miR-142-3p or cel-miR-67 as a control. Twenty-four hours after transfection, cells were starved with 0.5 % FBS for 30 min and exposed to different dose of diquat (0, 12.5, 25 µM), or etoposide (0, 25, 50uM), or cadmium (0, 5, 25 µM) as genotoxic agents for 24 hours. The cell viability against diquat were measured by MTS assay and the level of ROS after different dose of diquat treatment were measured and analysed by FACS. The cell viability against etoposide in miR-142-3p transfected IMR90 cells was measured by 5-chloromethylfluoresceindiacetate (CMFDA) staining in flow cytometry. Briefly, cell samples were stained with CMFDA (Molecular Probes, Eugene, OR) at a final concentration of 5 µM in PBS for 5 min at room temperature. After washing, the cells were fixed in 4 % paraformaldehyde for 20 min at room temperature and analyzed by flow cytometry at 520 nm. The cell viability against cadmium in miR-142-3p transfected IMR90 cells was measured by using trypan blue staining.

### Polyethylenimine (PEI) complexation of miR-142-3p

Polyehtylenimine (PEI) F25/miRNA complexes were prepared essentially as described previously for siRNAs^64^. Briefly, 40 μg miRNA was dissolved in 75 μl 10 mM HEPES / 150 mM NaCl, pH 7.4, and incubated for 10 min. 10 μl PEI F25-LMW (5 μg/μl) was dissolved in 75 μl of the same buffer, and pipetted to the miRNA solution after 10 min.

### Histopathology analysis

For hematoxylin-and-eosin (H&E) staining, tissue samples were fixed in 10 % neutral buffered formalin. Fixed tissues were embedded in paraffin and sectioned into slices of 5 μM each. Slides were stained in hematoxylin for 6 minutes, rinsed with water, and stained with eosin for another 1–2 minutes. Sections were dehydrated in 50 %, 70 %, 80 %, 95 %, and 100 % alcohol solutions, cleared with xylene, and mounted with a cover slip onto a labeled glass slide.

### OGTT and ipITT

For oral glucose tolerance test (OGTT) of all mice from young, old and high-fat diet model mice, after miRNA injection, mice were fasted overnight and orally administrated with 2 g/kg body weight of dextrose for oral glucose tolerance test (OGTT). For intraperitoneal insulin tolerance test (ipITT), mice were fated overnight after miRNA injection for 6 days and injected with 0.5 U/kg body weight of insulin (Novolin R insulin, Novolin) intraperitoneally. The level of glucose in blood was measured at fasting and at 15, 30, 60 and 120 min after administrating dextrose or insulin.

### Metabolomics

Liver samples were pulverized, weighed, and extracted with 8 times volume of extraction solution (5 mM ammonium acetate in methanol, containing 5nmol of U13C citrate/50mg, and 1nmol of U13C succinate/50mg as internal standards) to liver weight. The samples were homogenized twice with bead beat for 1min. Homogenized samples were subjected to froze-thaw (thaw with sonication for 5 min) cycle for three times. The samples were then centrifuged for 10 min. 20 µL of the extraction from each sample was mixed with 10 µL of internal standard working solution (including nine stable isotope-labeled acylcarnitines, lyso PC a C9:0, PC aa C28:0, PC aa C40:0 and SM C6:0), and then was extracted with 300 µL of extraction solvent (5 mM ammonium acetate in methanol). After vortexing, samples were kept at -20°C for 10 min, and then were centrifuged at 12,000 rpm for 10 min at 4°C. 150 µL of the extraction was diluted with 400 µL of mobile phase (methanol / water 97:3 with 10 mM ammonium acetate). 20 µL of diluted sample was injected into an electrospray ionization source and analyzed by Waters Xevo TQ MS (Waters, Milford, MA, USA). After taking 20ul for LC-MS, the reminder sample was added ¼ volumes of water and GC-MS internal standards and homogenized again for 1 min. The samples were centrifuged for another 10 min. The supernatant was dried under gentle nitrogen flow and derivatized with a two-step derivatization procedure. First, the samples were methoximized with 50 µl of methoxyamine hydrochloride (MOA, 15 mg/mL in pridine) at 30 °C for 90 min. The silylation step was done with 80 µL of N,O-Bis(trimethylsilyl) trifluoroacetamide (BSTFA, containing 1% TMCS) at 70 °C for 60 min. 1 QC sample was run three times during the analysis. The samples were analyzed by gas chromatography time-of-flight mass spectrometry (GC-TOFMS premier, Waters, USA). Separation was performed with a DB-5MS column. Helium was used as carrier gas at a consistent flow of 1mL/min. The oven program was as the following: started at 60 °C for 1min, 10 °C /min to 320 min and kept for 3 min. For some metabolites with high concentration in the urine such as creatinine and citrate, the samples were injected with 2 split ratios. One injected 1uL of sample with split ratio of 5, the other injected 0.5 µL of sample with split ratio of 100. Raw data was analyzed in Genedata Expressionist (Genedata, Basel, Switzerland) software. Metabolites annotation was performed with comparing the mass spectrum and retention time to commercially available libraries such as Fiehn library, and NIST library (GC-FS). QC sample was injected six times for coefficient of variation (CV) calculation for data quality control. Metabolites with CVs lower than 20 % were treated as accurate quantification, CVs between 20 % and 30 % were treated as relatively accurate quantification, while CVs higher than 30 % were treated as highly doubting results.

## Acknowledgments

Research reported in this publication was supported by the Albert Einstein Cancer Center Support Grant of the National Institutes of Health under award number P30CA013330.

Data published using the PerkinElmer P250 High-Capacity Slide Scanner must acknowledge shared instrumentation grant SIG #1S10OD019961-01.

## Author Contributions

H.J.J., B.M., J.C., C.T, X.F.W., XZ.W., J.Y. and Q.G. conducted the experiments and analyzed the results. H.J.J and X.F.W did the cell experiments. B.M. and J.Y conducted the analysis of RNA-sequencing. H.J.J and X.Z.W conducted the injection of miRNA mimics to mice. J.C. conducted GTT/ITT and physiological tests of mice under high-fat diet and aged mice. C.T. conducted the western blot of B-cells in AJ population. X.F.W, H.J.J., G.A., N.B., R.D.P, and Y.S. wrote the paper with input and approval of all authors.

## Data availability

All raw sequencing data were deposited in the NCBI Short Read Archive (SRA) under project numbers PRJNA940421 (microRNA-seq of B cells from centenarians and controls) and PRJNA940672 (RNA-seq of MCF7).

