## Extended Data Figures for "The regulation of Insulin/IGF-1 signaling by miR-142-3p associated with human longevity"

**Extended Data Fig. 1 qRT-PCR validation of differentially expressed miRNAs in centenarians and controls.**

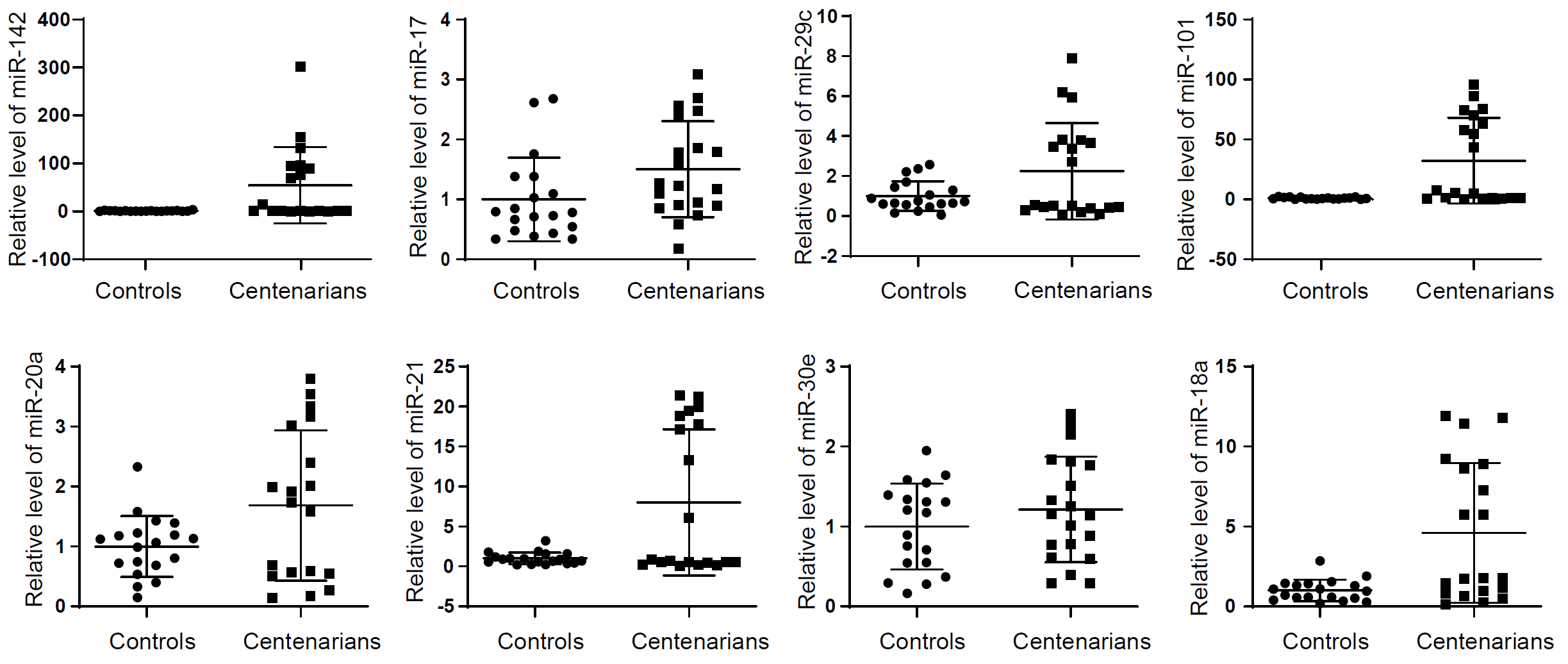

**Extended Data Fig. 2 Effect of miR-142-3p overexpression on IIS signaling pathway.** (a) The protein level of IIS molecules after miR-142-3p transfection into IMR90 cells. (b) The quantification analysis for the protein level of IIS molecules from (a). (c) Relative expression level of miR-142-3p in offsprings of AJ centenarians and age-matched controls in different age groups. (d) vocalno plot of differentially expressed genes between miR-142-3p-transfected MCF7 cells and controls (False Discovery Rate < 0.05, fold change > 1.5). Colored dots correspond to miRNAs significanly enriched in centenarians (red dots), or significantly enriche in elderly controls (blue), or not significant in either groups (grey). (e) Dot plots representing the changes in GSEA hallmark pathways after overexpression of miR-142-3p in MCF7 cells. (f) Venn diagram summarizing the overlap between downregulated genes from RNA-Seq (blue circle) and predicted targets of miR-142-3p from Targetscan (grey circle) and MicroT-CDS (orange circle). (g) qRT-PCR validation of differentially expressed genes involving IIS pathway and predicted as miR-142-3p targets. Data are expressed as mean ± SEM. *p < 0.05; **p <0.01; ***p < 0.001. (h) Protein level of IGF1R and FURIN in miR-142-3p overexpressed MCF7 cells.

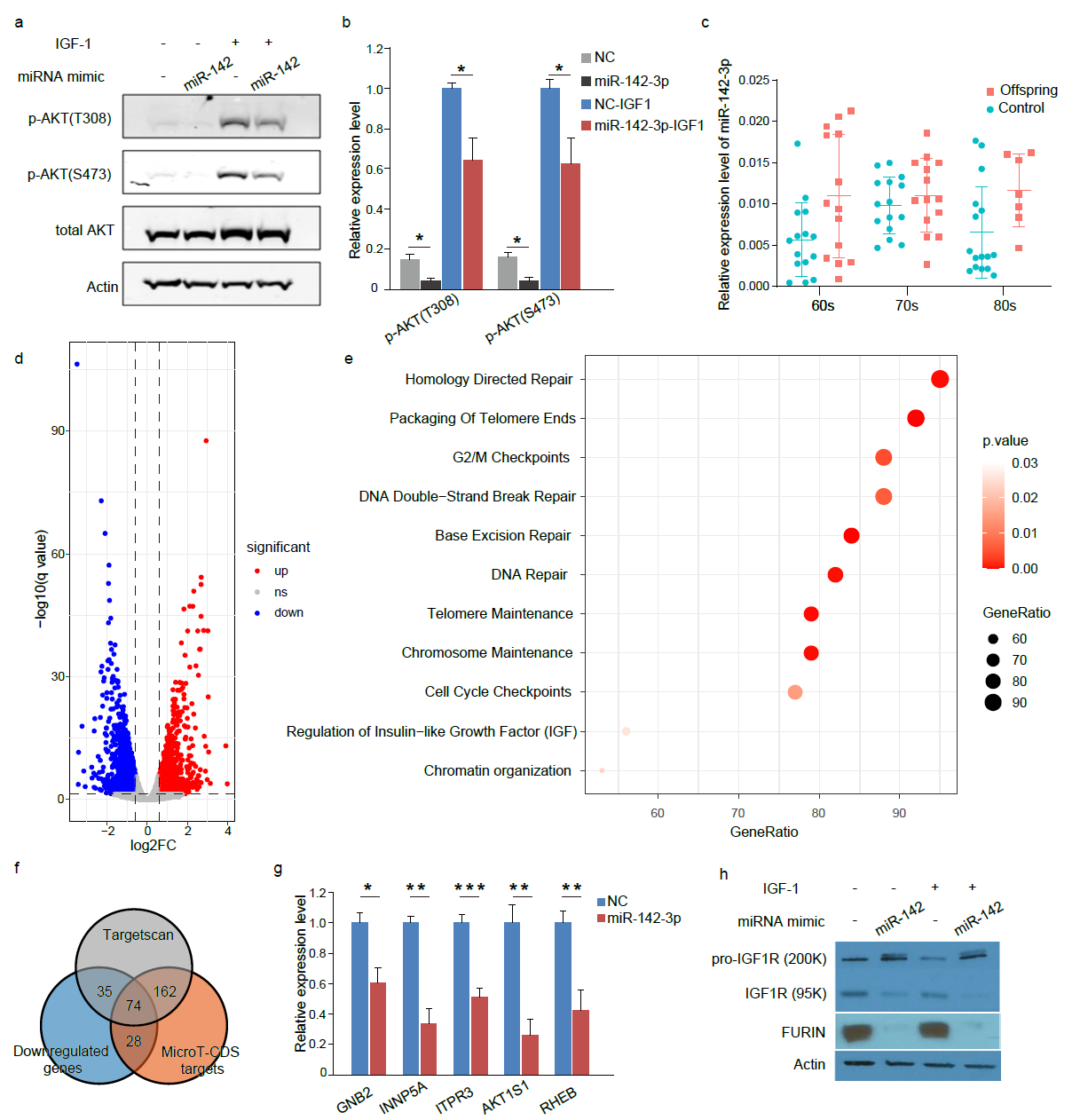

**Supplementary Figure 3. Protective effects of miR-142-3p on human normal fibroblast IMR90 cells against genotoxic stressors.** (A) Cell viability (%) after etoposide treatment (25 and 50uM) for 24 hours in miR-142-3p transfected IMR90 cells by using FACS analysis. (B) Cell viability (%) after cadmium treatment (5 and 25 uM) for 24 hours in miR-142-3p transfected IMR90 cells by using trypan blue staining.

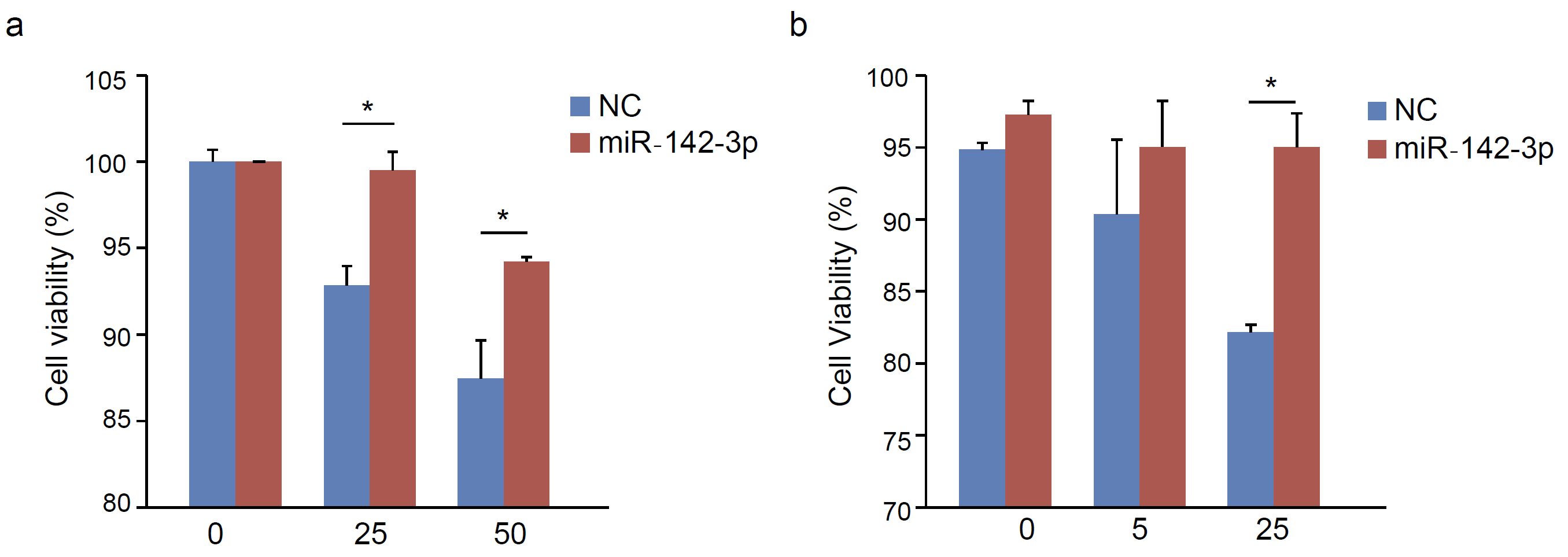

**Supplementary Figure 4. Uptake of PEI-complexed miR-142-3p in different mouse tissues**

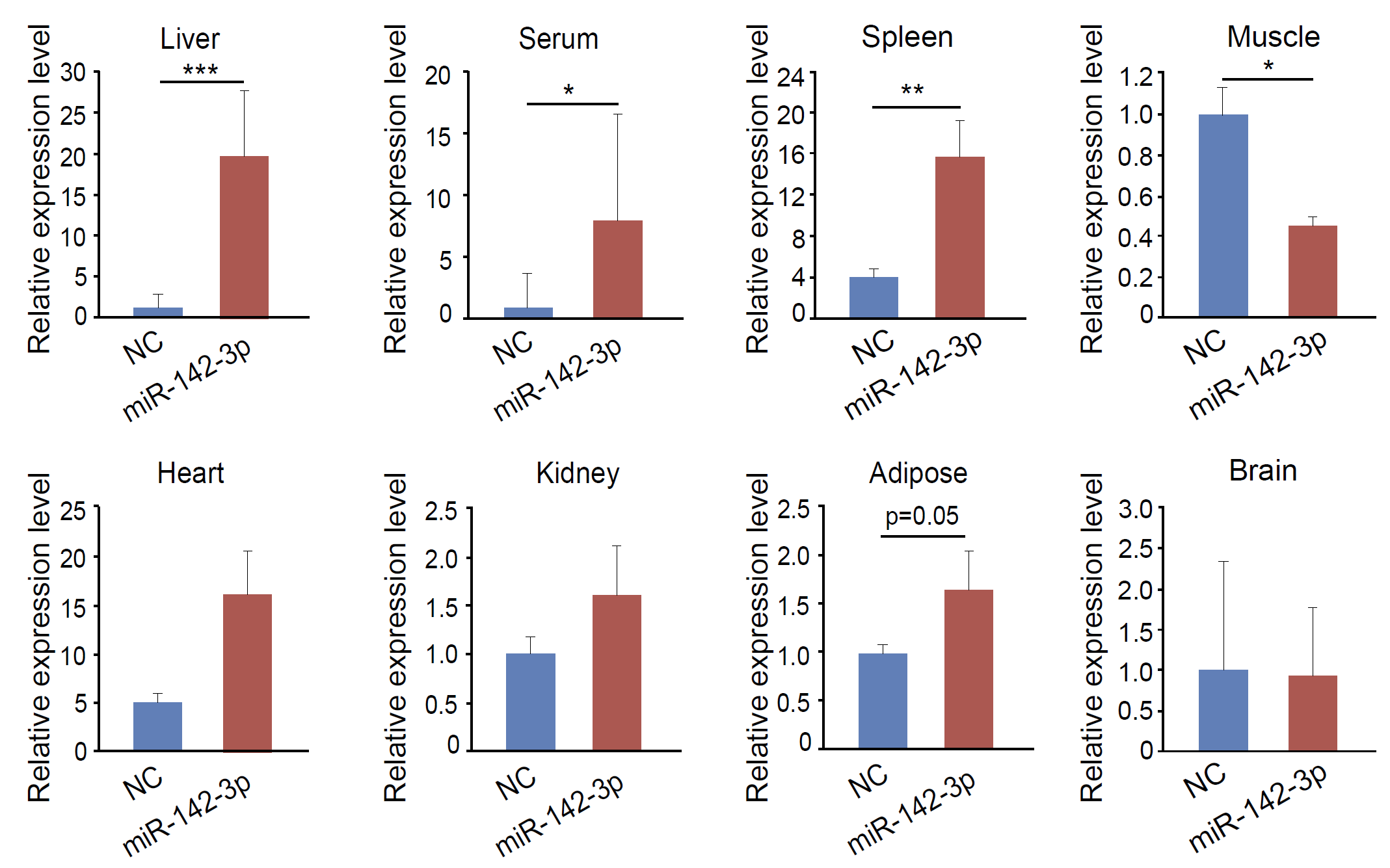

**Supplementary Figure 5. physiological responses of miR-142-3p-injected mice under HFD treatment and metabolic alterations upon miR-142-3p overexpression in liver tissue.** (a) Percentage of weight gain of miR-142-3p-injected mice and control-injected mice in 4 weeks of HFD feeding. (b) Average day and night Respiratory exchange ratio (REF) of miR-142-3p-injected mice and control-injected mice. (c) Average day and night energy expenditure values of miR-142-3p-injected mice and control-injected mice. (d-e) Average day and night physical activity levels of miR-142-3p-injected mice and control-injected mice. (f-g) Validated model plots of GC/MS (f) and LC/MS (g) obtained by permutation test.

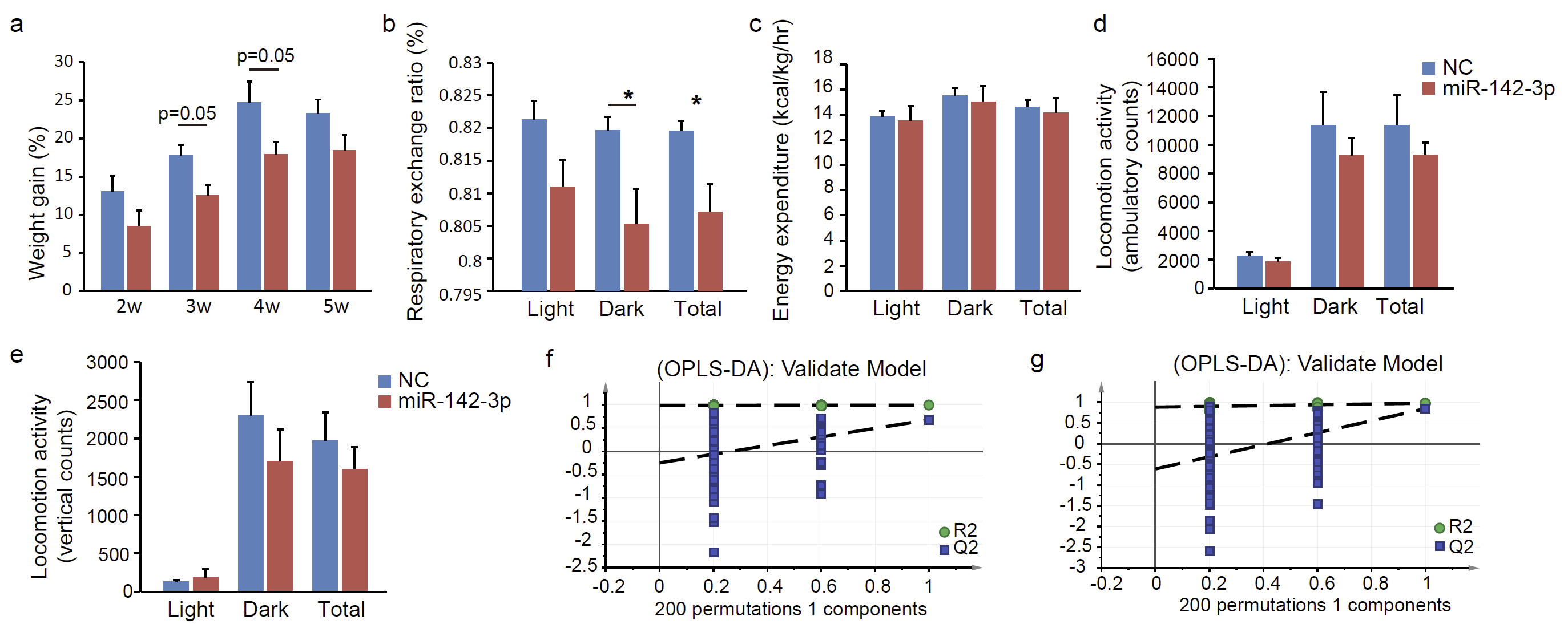
