## Supplementary Tables for "The regulation of Insulin/IGF-1 signaling by miR-142-3p associated with human longevity"

**Supplementary Table 1. Differentially expressed miRNAs in B cells between centenarians (n=20) and controls (n=20),** **FDR < 0.05, Fold Change > 2**

| **miRNA** | **Mean-Control** | **Mean-Centenarian** | **FDR** | **Fold Change** |
| --- | --- | --- | --- | --- |
| hsa-miR-142-5p | 981.79 | 42584.27 | 7.42E-18 | 43.37 |
| hsa-miR-142-3p | 231.47 | 7476.27 | 1.56E-15 | 32.30 |
| hsa-miR-10b-5p | 51.62 | 1456.43 | 1.65E-12 | 28.24 |
| hsa-miR-301a-3p | 14.39 | 387.66 | 3.58E-13 | 27.01 |
| hsa-miR-19b-3p | 96.99 | 2260.13 | 8.80E-12 | 23.32 |
| hsa-miR-29b-3p | 30.18 | 575.24 | 4.61E-13 | 19.07 |
| hsa-miR-101-3p | 43.71 | 763.46 | 1.56E-15 | 17.47 |
| hsa-miR-301b | 15.26 | 183.54 | 3.45E-09 | 12.05 |
| hsa-miR-29c-3p | 100.79 | 1052.76 | 1.17E-10 | 10.45 |
| hsa-miR-21-5p | 4061.25 | 38044.90 | 6.71E-11 | 9.37 |
| hsa-miR-424-5p | 12.03 | 105.35 | 1.19E-05 | 8.75 |
| hsa-miR-33a-5p | 6.70 | 57.38 | 8.44E-09 | 8.53 |
| hsa-miR-660-5p | 21.10 | 127.91 | 1.15E-08 | 6.07 |
| hsa-miR-18a-5p | 98.91 | 595.30 | 9.61E-09 | 6.02 |
| hsa-miR-148a-3p | 6915.40 | 39795.65 | 1.30E-08 | 5.75 |
| hsa-miR-30d-3p | 21.44 | 108.10 | 6.75E-08 | 5.03 |
| hsa-miR-192-5p | 409.14 | 3434.09 | 5.32E-04 | 4.87 |
| hsa-miR-30a-5p | 142.96 | 695.59 | 5.44E-05 | 4.87 |
| hsa-miR-33a-3p | 7.32 | 28.27 | 3.98E-05 | 3.84 |
| hsa-miR-29a-5p | 12.39 | 47.32 | 2.09E-04 | 3.83 |
| hsa-miR-141-3p | 80.54 | 300.55 | 1.43E-05 | 3.73 |
| hsa-miR-7-5p | 28.44 | 104.21 | 6.36E-04 | 3.67 |
| hsa-miR-7-1-3p | 18.10 | 64.84 | 3.40E-05 | 3.58 |
| hsa-miR-27a-5p | 12.66 | 44.61 | 2.06E-03 | 3.51 |
| hsa-miR-374b-5p | 19.89 | 66.69 | 9.76E-05 | 3.35 |
| hsa-miR-30b-5p | 786.45 | 2588.36 | 2.30E-05 | 3.29 |
| hsa-miR-454-3p | 110.73 | 353.73 | 1.39E-04 | 3.20 |
| hsa-miR-486-3p | 8.95 | 28.69 | 3.73E-04 | 3.17 |
| hsa-miR-30e-5p | 6207.33 | 19652.84 | 3.37E-05 | 3.17 |
| hsa-miR-181a-3p | 106.60 | 334.51 | 4.76E-04 | 3.14 |
| hsa-miR-20a-5p | 963.73 | 2998.27 | 2.00E-05 | 3.11 |
| hsa-miR-335-3p | 24.44 | 74.45 | 7.77E-03 | 3.04 |
| hsa-miR-223-3p | 79.94 | 239.68 | 1.30E-04 | 3.00 |
| hsa-let-7g-3p | 9.00 | 26.41 | 4.22E-05 | 2.92 |
| hsa-miR-27a-3p | 759.92 | 2192.59 | 1.43E-04 | 2.89 |
| hsa-miR-34a-3p | 9.35 | 26.72 | 2.47E-03 | 2.85 |
| hsa-miR-4518 | 7.03 | 20.26 | 4.88E-05 | 2.84 |
| hsa-miR-3681-5p | 41.12 | 104.94 | 1.03E-02 | 2.55 |
| hsa-miR-576-5p | 28.71 | 72.25 | 1.62E-03 | 2.52 |
| hsa-miR-16-2-3p | 130.08 | 319.90 | 2.15E-04 | 2.46 |
| hsa-miR-106b-5p | 247.16 | 596.30 | 3.38E-05 | 2.41 |
| hsa-miR-15a-5p | 288.71 | 696.10 | 1.62E-04 | 2.41 |
| hsa-miR-148b-5p | 64.98 | 149.29 | 7.75E-04 | 2.30 |
| hsa-miR-15b-3p | 118.29 | 240.49 | 9.43E-04 | 2.03 |
| hsa-miR-125a-5p | 692.59 | 292.62 | 1.62E-03 | 0.42 |
| hsa-miR-6087 | 18.53 | 11.45 | 7.77E-03 | 0.35 |
| hsa-miR-125b-2-3p | 34.83 | 9.63 | 1.28E-02 | 0.28 |
| hsa-let-7c-5p | 474.69 | 115.56 | 1.93E-14 | 0.24 |
| hsa-miR-125b-5p | 159.08 | 31.57 | 7.68E-06 | 0.20 |

**Supplementary Table 2. The overlap miRNAs between lymphocytes and plasma**

| miRNAs | Fold Change | |
| --- | --- | --- |
|  | LCLs | Plasma |
| miR-142-3p | 32.3 | 10.84 |
| miR-101-3p | 17.47 | 5.83 |
| miR-301b | 12.05 | 8.34 |
| miR-29c-3p | 10.45 | 5.64 |
| miR-21-5p | 9.37 | 5.05 |
| miR-660-5p | 6.07 | 7.47 |
| miR-148a-3p | 5.75 | 5.51 |
| miR-27a-3p | 2.89 | 5.95 |
| miR-15a-5p | 2.41 | 28.45 |

**Supplementary Table 3. Differentially expressed genes between miR-142-3p overexpressed MCF7 cells and control MCF7 cells FDR<0.05, Fold Change>1.5**

| **Gene** | **log2FC** | **padj** | **Gene** | **log2FC** | **padj** | **Gene** | **log2FC** | **padj** |
| --- | --- | --- | --- | --- | --- | --- | --- | --- |
| IL6ST | -2.29 | 1.10E-73 | ZNF532 | -1.80 | 1.11E-10 | FAM114A1 | -0.79 | 1.99E-06 |
| AKT1S1 | -1.93 | 1.71E-53 | COPS7A | -0.94 | 1.77E-10 | CXADR | -1.23 | 2.72E-06 |
| CTTN | -1.66 | 3.08E-36 | PGM1 | -1.07 | 1.97E-10 | ZNF618 | -0.98 | 2.90E-06 |
| TWF1 | -1.50 | 1.62E-32 | KIF5B | -0.84 | 4.01E-10 | FBXO21 | -0.59 | 3.30E-06 |
| SLC37A3 | -1.74 | 2.85E-30 | FMNL2 | -1.30 | 5.79E-10 | RARG | -0.61 | 4.16E-06 |
| RAB12 | -1.56 | 7.69E-30 | ITPR3 | -1.45 | 6.63E-10 | RHEB | -0.81 | 4.53E-06 |
| VAMP3 | -1.46 | 1.17E-29 | ITGAV | -0.80 | 7.78E-10 | DHX33 | -0.65 | 4.62E-06 |
| WASL | -1.44 | 3.45E-28 | MOB4 | -1.09 | 8.23E-10 | SKP2 | -0.76 | 4.71E-06 |
| RAB3B | -2.01 | 1.95E-25 | RAB1A | -0.85 | 8.35E-10 | MTCH1 | -0.71 | 6.12E-06 |
| TSEN34 | -1.51 | 2.47E-24 | SH2B1 | -0.82 | 1.17E-09 | ANKRD33B | -1.44 | 6.30E-06 |
| INPP5A | -1.19 | 2.03E-23 | CCNJ | -1.05 | 1.44E-09 | MUC5B | -2.42 | 6.56E-06 |
| PSMB5 | -1.52 | 2.57E-23 | MRFAP1 | -1.10 | 1.53E-09 | ZCCHC14 | -0.89 | 7.14E-06 |
| NT5DC2 | -1.26 | 2.96E-23 | MARCKS | -1.19 | 1.98E-09 | TXLNA | -0.66 | 7.95E-06 |
| CFL2 | -1.81 | 8.14E-23 | TMEM115 | -0.76 | 2.92E-09 | IPO7 | -0.59 | 8.19E-06 |
| MANBAL | -1.42 | 1.26E-21 | CLOCK | -0.80 | 4.91E-09 | TP53INP2 | -0.85 | 1.96E-05 |
| QKI | -1.08 | 3.97E-21 | GOLGA1 | -1.00 | 7.09E-09 | WDR62 | -0.66 | 2.62E-05 |
| GNB2 | -1.04 | 5.67E-20 | EML4 | -0.87 | 9.26E-09 | CLTA | -0.69 | 2.81E-05 |
| LRRC59 | -1.04 | 3.40E-17 | RAC1 | -0.83 | 1.10E-08 | DNAJB5 | -0.75 | 3.32E-05 |
| TGFBR1 | -1.37 | 5.75E-17 | KLF13 | -1.19 | 1.63E-08 | SCUBE3 | -1.37 | 3.46E-05 |
| PPP1R37 | -1.23 | 8.99E-17 | MLXIP | -1.19 | 2.10E-08 | SEMA3D | -0.86 | 3.58E-05 |
| MORF4L2 | -1.05 | 1.26E-16 | SLC25A22 | -0.71 | 2.59E-08 | SYPL1 | -0.61 | 3.66E-05 |
| THSD4 | -1.41 | 1.95E-16 | RGL2 | -0.83 | 2.80E-08 | BIRC3 | -1.76 | 4.28E-05 |
| LOXL1 | -1.27 | 3.41E-16 | STX12 | -0.84 | 3.33E-08 | RNF157 | -0.85 | 4.59E-05 |
| ANKFY1 | -0.98 | 1.85E-15 | TMTC3 | -0.81 | 5.01E-08 | CRK | -0.64 | 5.18E-05 |
| CASK | -0.95 | 7.55E-15 | DYNC1LI2 | -0.74 | 5.93E-08 | RBMS1 | -0.61 | 5.74E-05 |
| NR1D2 | -1.03 | 1.57E-14 | TMED7 | -0.91 | 7.88E-08 | MTMR9 | -0.62 | 8.20E-05 |
| EPN1 | -1.25 | 1.66E-14 | CNIH4 | -1.08 | 8.32E-08 | CLDN12 | -0.79 | 9.26E-05 |
| BOD1 | -1.15 | 1.76E-14 | TGFB2 | -1.02 | 1.11E-07 | ANKS1A | -0.78 | 1.48E-04 |
| HSPA1B | -0.97 | 2.14E-14 | RIPPLY3 | -1.44 | 1.17E-07 | RAB40C | -0.58 | 1.66E-04 |
| TPM4 | -0.89 | 2.17E-14 | TFG | -0.69 | 1.17E-07 | TRIM14 | -0.67 | 2.12E-04 |
| RAB3D | -0.99 | 2.63E-14 | WIZ | -1.21 | 1.39E-07 | SNX18 | -0.60 | 2.13E-04 |
| CLIC4 | -1.26 | 2.87E-14 | RPRD1A | -0.76 | 2.50E-07 | SERAC1 | -0.95 | 6.47E-04 |
| PAFAH1B2 | -1.18 | 5.37E-14 | TAB2 | -0.69 | 2.99E-07 | WHAMM | -0.65 | 7.82E-04 |
| CHMP3 | -0.90 | 1.34E-13 | MAP3K11 | -1.04 | 3.37E-07 | FOXO1 | -0.61 | 8.44E-04 |
| NR2F6 | -1.37 | 2.64E-13 | RAB3A | -0.79 | 4.53E-07 | SLC17A5 | -0.68 | 1.35E-03 |
| COG4 | -0.88 | 2.93E-13 | ASH1L | -0.98 | 4.88E-07 | FBXO3 | -0.62 | 1.45E-03 |
| LLGL2 | -0.86 | 4.46E-13 | ITGB8 | -0.91 | 5.05E-07 | GPR137C | -0.81 | 1.51E-03 |
| CDC14A | -1.22 | 5.90E-13 | ATP1B1 | -0.73 | 5.68E-07 | SPRED1 | -0.59 | 1.87E-03 |
| ZCCHC24 | -1.06 | 8.22E-13 | PES1 | -0.67 | 5.70E-07 | MBD6 | -0.59 | 4.03E-03 |
| INHBA | -1.28 | 2.59E-12 | STAU1 | -0.63 | 7.71E-07 | ROCK2 | -0.71 | 7.16E-03 |
| HMGA1 | -1.03 | 3.06E-11 | PCGF3 | -0.62 | 8.98E-07 | PTPN23 | -0.80 | 7.41E-03 |
| LATS1 | -0.84 | 3.93E-11 | NUCKS1 | -0.69 | 9.84E-07 | TNFRSF13C | -1.44 | 7.82E-03 |
| MXRA7 | -1.24 | 4.60E-11 | MYBL1 | -0.74 | 1.19E-06 | ERG | -1.22 | 1.70E-02 |
| SLC30A9 | -0.86 | 7.26E-11 | FNTB | -0.82 | 1.38E-06 | MMD | -0.80 | 3.02E-02 |
| HGS | -0.91 | 1.04E-10 | DCUN1D4 | -0.88 | 1.47E-06 | ZNF462 | -0.66 | 4.70E-02 |
| HECTD1 | -0.73 | 1.11E-10 | ATG16L1 | -0.64 | 1.70E-06 |  |  |  |

**Supplementary Table 4. Differentially expressed metabolites (GC-MS) between mice injected with PEI-complexed miR-142-3p and control mice injected with PEI-complexed cel-miR-67. *p* < 0.05, VIP > 1**

| **Metabolites** | **Control** | **miR-142-3p** | **T test** | **VIP** | **Fold change** |
| --- | --- | --- | --- | --- | --- |
| hypotaurine | 34.60 | 133.96 | 0.05 | 6.86 | 3.87 |
| 2-aminobutyric acid | 13.15 | 28.87 | 0.00 | 3.14 | 2.20 |
| Beta- alanine | 8.59 | 17.38 | 0.01 | 2.29 | 2.02 |
| DL-Ornithine | 9.07 | 18.13 | 0.02 | 2.20 | 2.00 |
| L-lysine | 12.10 | 20.41 | 0.01 | 2.27 | 1.69 |
| L-threonine | 98.44 | 163.63 | 0.00 | 6.58 | 1.66 |
| putrescine | 21.68 | 32.46 | 0.00 | 2.64 | 1.50 |
| L-valine | 135.20 | 181.99 | 0.02 | 5.11 | 1.35 |
| d-Erythrotetrofuranose | 34.00 | 45.49 | 0.04 | 2.33 | 1.34 |
| Isoleucine | 57.33 | 75.56 | 0.05 | 2.96 | 1.32 |
| Glycerol 3-phosphate | 424.34 | 388.42 | 0.02 | 4.58 | 0.92 |
| Beta-glycerolphosphate | 13.96 | 11.46 | 0.05 | 1.13 | 0.82 |
| 2-hydroxypentanedioic acid | 10.14 | 7.93 | 0.02 | 1.15 | 0.78 |
| Nicotinamide | 42.73 | 30.25 | 0.00 | 2.95 | 0.71 |
| Cytindine-5'-monophosphate | 24.07 | 14.42 | 0.02 | 2.28 | 0.60 |
| 2-aminoethanethiol | 6.54 | 1.44 | 0.00 | 1.95 | 0.22 |

**Supplementary Table 5. Differentially expressed metabolites (LC-MS) between mice injected with PEI-complexed miR-142-3p and control mice injected with PEI-complexed cel-miR-67. *p* < 0.05, VIP > 1**

| **Metabolites** | **Controls** | **miR-142-3p** | **t TEST** | **VIP** | **Fold change** |
| --- | --- | --- | --- | --- | --- |
| PC ae C38:2 | 25.90 | 43.22 | 0.02 | 1.93 | 1.67 |
| PC ae C36:3 | 10.74 | 15.61 | 0.00 | 1.10 | 1.45 |
| PC ae C34:2 | 15.94 | 23.13 | 0.00 | 1.37 | 1.45 |
| PC ae C36:2 | 37.71 | 51.52 | 0.01 | 1.76 | 1.37 |
| lysoPC a C18:2 | 33.65 | 41.42 | 0.00 | 1.42 | 1.23 |
| PC ae C36:1 | 42.16 | 50.88 | 0.00 | 1.46 | 1.21 |
| SM C24:0 | 83.52 | 100.32 | 0.02 | 2.07 | 1.20 |
| SM C16:0 | 101.20 | 90.88 | 0.04 | 1.38 | 0.90 |
| PC aa C40:6 | 361.36 | 312.96 | 0.05 | 3.16 | 0.87 |
| PC aa C38:6 | 1278.40 | 1088.00 | 0.01 | 6.75 | 0.85 |
